# Short-term memory errors are strongly associated with a drift in neural activity in the posterior parietal cortex

**DOI:** 10.1101/2024.09.03.610917

**Authors:** Joon Ho Choi, Sungwon Bae, Jiho Park, Minsu Yoo, Chul-Hoon Kim, L. Ian Schmitt, Ji-Woong Choi, Jong-Cheol Rah

**Author notes:** These authors contributed equally (Joon Ho Choi, Sungwon Bae). Corresponding authors: Joon Ho Choi, Sensory and Motor Systems Neuroscience Group, Korea Brain Research Institute, 61, Cheomdan-ro, Dong-Gu, Daegu 41062, Korea; Jong-Cheol Rah, Sensory and Motor Systems Neuroscience Group, Korea Brain Research Institute, 61, Cheomdan-ro, Dong-Gu, Daegu 41062, Korea.

## Abstract

Understanding the neural mechanisms behind short-term memory (STM) errors is crucial for unraveling cognitive processes and addressing deficits associated with neuropsychiatric disorders. This study examines whether STM errors arise from misrepresentation of sensory information or decay in these representations over time. Using 2-photon calcium imaging in the posterior parietal cortex (PPC) of mice performing a delayed match-to-sample task, we identified a subset of PPC neurons exhibiting both directional and temporal selectivity. Contrary to the hypothesis that STM errors primarily stem from mis-encoding during the sample phase, our findings reveal that these errors are more strongly associated with a drift in neural activity during the delay period. This drift leads to a gradual divergence away from the correct representation, ultimately leading to incorrect behavioral responses. These results emphasize the importance of maintaining stable neural representations in the PPC for accurate STM. Furthermore, they highlight the potential for therapeutic interventions aimed at stabilizing PPC activity during delay periods as a strategy for mitigating cognitive impairments in conditions like schizophrenia.

## Introduction

Short-term memory (STM) refers to the capacity to retain immediately relevant information over brief periods, typically on the order of seconds to minutes. While STM is integral to numerous cognitive functions, its representations are fragile and prone to errors, often leading to suboptimal behavior. Despite the significant behavioral consequences of STM errors, there is currently no circuit-level understanding of their underlying mechanisms, primarily due to a lack of systematic investigation. Gaining insight into the circuit mechanisms behind STM errors is critical for understanding information retention and its implications for psychiatric disorders, including schizophrenia and attention deficit hyperactivity disorder, which are frequently associated with STM deficits^1–3^. This study aims to investigate the circuit-level basis of STM errors by examining two potential mechanisms that may underlie their generation: (1) Inadequate encoding of information in neural activity during the initial presentation of stimuli, and (2) Disruption of encoded information during the delay period, leading to retention failures.

Along with the prefrontal cortex (PFC), the posterior parietal cortex (PPC) is a critical brain region involved in maintaining information during STM^4–6^. We chose to investigate the activity of the PPC for several reasons. As a major multimodal association cortex, the PPC is ideally positioned to integrate sensory inputs from multiple modalities and relay this accumulated information to prefrontal areas^6,7^. This makes the PPC a plausible origin for erroneous activity during STM error trials, especially if inadequate evidence accumulation or misperception is reflected in both the PFC and PPC. In both primates and rodents, a subset of PPC neurons demonstrates direction-predictive firing rate changes in a graded manner, encoding the clarity of evidence during decision-making tasks such as the random dot motion discrimination task or the Poisson click task^8,9^. In contrast, neurons in prefrontal motor areas show more categorical activity patterns, suggesting that cumulative evidence in the PPC is transformed into discrete decisions within the PFC. Based on these findings, we hypothesized that graded degradation of encoded information, which could lead to STM errors, would be more readily observable in the PPC than in the PFC. By focusing on the PPC, we aimed to capture subtle changes in sensory encoding and retention that might underlie STM errors.

Despite substantial evidence implicating the PPC as a critical substrate for STM, the mechanisms underlying information maintenance in this circuit remain incompletely understood. In a previous study, mice were trained to navigate a virtual T-maze by choosing a left or right turn based on visual cues presented during the initial phase of each trial, while PPC activity was recorded. Since the visual cue was available only during the first half of the trial, successful navigation required the mice to rely on STM. This study demonstrated that PPC activity involves a complex population encoding, characterized by direction- and phase-selective sequences across multiple neurons^6^. However, interpreting PPC activity during such tasks as a direct reflection of visual STM is challenging due to its navigational attributes^10–12^. For instance, Krumin et al.^13^ demonstrated that sequential PPC activity during a similar behavioral task could be accurately predicted by the combination of the animal’s heading direction and spatial location within the virtual corridor. Further evidence of task^13^-relevant PPC activity comes from mice performing a visually guided delayed go/no-go task^6,14^. In this paradigm, the PPC exhibited distinct activity patterns, with a substantial number of neurons selectively encoding ‘go’ signals, while ‘no-go’ signals were represented more weakly^6,14^. This observation raises important questions about whether these activities genuinely reflect the retained visual signal or are more related to the suppression and initiation of impulsive movement^15^.

To more directly investigate how sensory information is retained during the delay period and how disruptions in this process contribute to errors, we recorded PPC activity while mice performed a delayed match-to-sample task that did not involve navigation. A substantial fraction of imaged PPC neurons exhibited activity patterns associated with memory-guided decision-making. Notably, these patterns closely resembled those observed in previous STM studies^6^, supporting the idea of a shared mechanism for encoding short-term representations across different tasks. A comparison of PPC activity between correct and incorrect trials revealed key differences in the neural dynamics underlying STM errors. Specifically, errors were associated with weak initial encoding of the sensory signal, followed by spontaneous drift of neural activity toward an incorrect representation during the delay period. These findings shed light on the circuit mechanisms underlying STM errors, providing critical insight into the processes of sensory encoding and maintenance. Additionally, this work opens avenues for future research into how STM mechanisms may be disrupted in neuropsychiatric disorders.

## Results

### Sequence of choice-dependent neuronal activity in PPC

Our objective was to determine the extent to which PPC activity is critical for maintaining sensory information during an STM-dependent perceptual decision-making task, excluding confounding factors such as spatial location and head direction. To this end, we trained seven transgenic mice expressing GCaMP6f (C57BL/6J-Tg(Thy1-GCaMP6f)GP5.5Dkim/J) to perform STM tasks. In this task, head-restrained mice were presented with moving gratings displayed on a screen in front of them, with the direction of motion (leftward or rightward) determined pseudo-randomly. After the visual stimulus, there was approximately a 2-s delay during which time no visual input was provided. Following this delay, a bifurcated lick port was introduced within the mice’s reach. Correct responses, consisting of licking the port corresponding to the motion direction of the grating, were rewarded with approximately 5 μL of tap water. Incorrect responses, defined as licking the opposite port, triggered an extended inter-trial interval of 2 s as a "time-out" penalty (see Fig. 1A). After extensive training, the mice demonstrated successful task performance, achieving a discrimination rate exceeding 70%. Importantly, the mice displayed no directional bias following training, indicating reliable task learning and unbiased decision-making (Fig. 1B-C).

**Figure 1:**
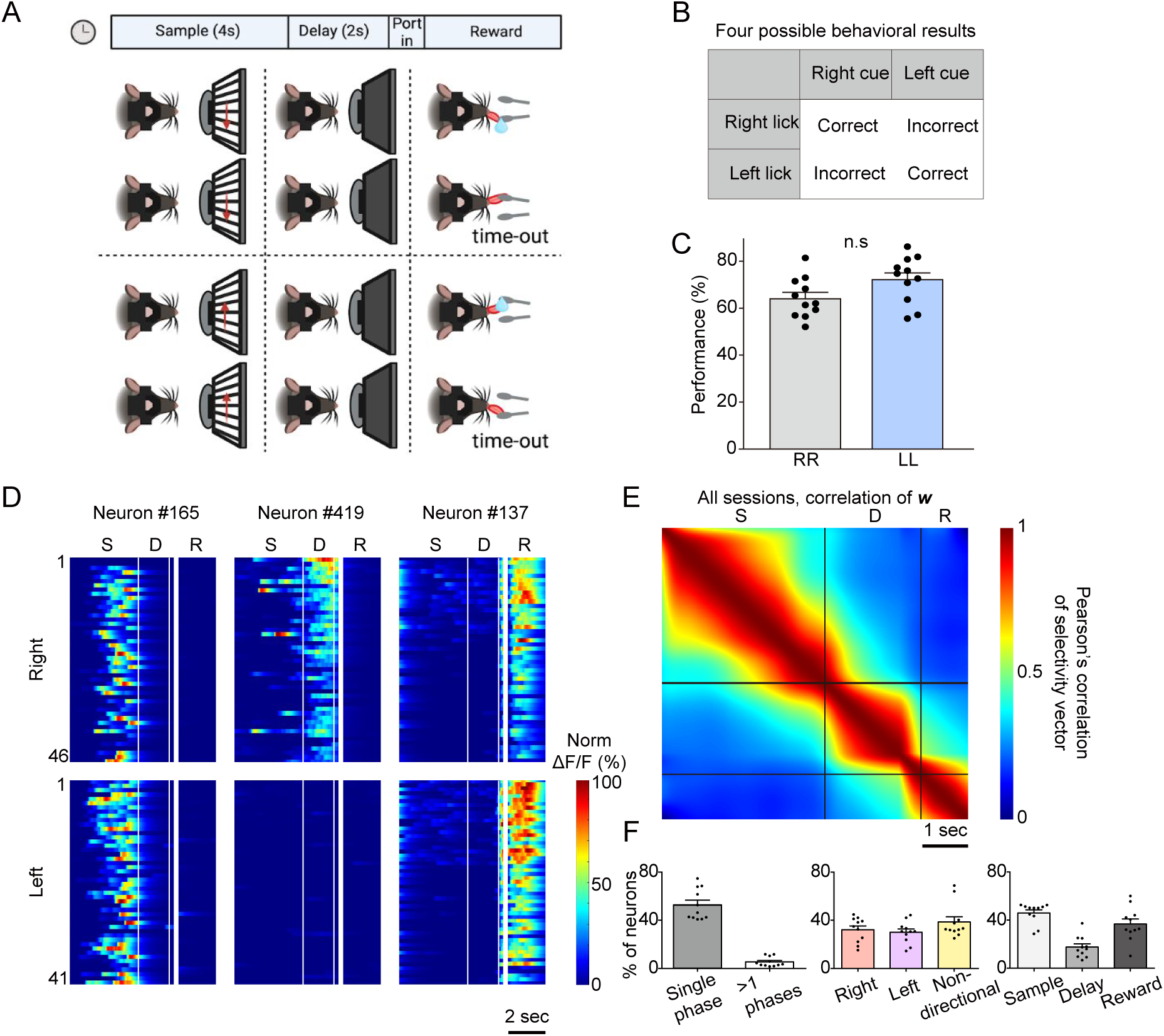
Ca^2+^ responses of PPC neurons during short-term memory task. **A.** The schematic illustrates the behavioral task designed to assess short-term memory in head-restrained mice. During the sample phase, black-and-white phase-reversing gratings moved either leftward or rightward (or downward or rightward) for 4 s on a screen in front of the mouse. After a 2-s delay, two laterally positioned lick ports were placed within reach of the mouse’s tongue. Correct licking at the corresponding lick port was rewarded with water, whereas incorrect licking resulted in a time-out as a punishment. **B.** Illustration of the four possible behavioral outcomes in the task. **C.** Performance comparison between rightward and leftward trials in well-trained mice. No significant difference was observed (two-tailed paired *t*-test; n=11, *p* < 0.05). Error bars represent the standard error of the mean (SEM) across sessions. n.s: not significant. **D.** Representative neuronal activity recorded during the behavioral tasks. Each row shows the color-coded normalized **Δ**F/F. **E.** The color-coded autocorrelation matrix of mean response vectors, ***w***, calculated from the coding direction (CD) of correct trials. ***w*** was calculated over moving 0.5-s time windows. Data represent 11 sessions, 6 mice, and 4756 total neurons S: sample phase, D: delay phase, and R: response phase. **F.** Proportion of neurons displaying task-dependent activity based on a General Linear Model (GLM). (Left) Fraction of neurons with single-phase selectivity, and neurons active in two or more phases as a percentage of all recorded neurons. (Middle) Percentages of single-phase selective neurons with and without directional selectivity. (Right) Phase selectivity distribution among single-phase selective neurons.

During task performance, we recorded the activity of PPC neurons by analyzing fluorescence changes in the GCaMP6f-expressing layer (L) 2/3 neurons using two-photon laser scanning microscopy. The identification of the PPC region was guided by stereotaxic coordinates based on mesoscopic connectivity analysis, which revealed distinct anterograde and retrograde connections compared to neighboring regions^16^. Active neurons were identified through post hoc analysis using the CaImAn software^17^. We observed a variety of neuronal responses within the PPC; however, a substantial proportion of neurons consistently exhibited selectivity based on direction, task phase, or both across trials (Fig. 1D). In addition to the striking consistency of these responses across trials, we computed the time-wise correlation of mean response vectors calculated every 0.5 s (15 image frames) across all recorded neurons. This analysis revealed that neural activity was consistent within behavioral phases, further supporting the robustness of PPC encoding during task performance (Fig. 1E; see Methods).

To objectively categorize active neurons based on their response patterns, we employed a modified generalized linear model (GLM), as described in the Methods section^18^. The time-correlation of activity vectors from the neurons sorted by our GLM was nearly identical to those from all recorded neurons (Supplementary Fig. 1), suggesting that the phase-dependent feature of the activity is well-preserved in the GLM-sorted neurons. Our analysis revealed that nearly half of the active neurons in the PPC exhibited selectivity for either phase or stimulus direction (*p* < 0.01), totaling 2,664 out of 4,569 neurons (Supplementary Fig. 2). Of these, approximately 60% (1,609/2,664 neurons; 873 for the right signal and 736 for the left signal) exhibited directional selectivity, while the remaining neurons responded similarly to both directions (Supplementary Fig. 3A-C). Among the 1,609 directionally selective neurons, approximately 41% displayed phase selectivity, with 661 neurons showing phase selectivity during the sample phase and 315 during the delay phase. Consistent with previous studies, we found that PPC neurons rarely exhibited sustained activity across multiple phases of the task^6^, with less than 8% (202/2664) of neurons demonstrating phase selectivity during both the sample and delay phases (Fig. 1F and Supplementary Fig. 4).

When the neurons were sorted based on the onset of peak responses, a clear directional-selective sequence of neuronal activity emerged, closely resembling the findings of Harvey et al.^6^ (Fig. 2A). This result underscores the replicability of the directional-selective neural sequence in the PPC, even within a behavioral paradigm that excluded spatial navigation. To assess the predictive capabilities of the GLM-selected neurons, we verified the direction selectivity of each neuron using receiver operating characteristic (ROC) analysis (Fig. 2C-D). While ROC analysis identifies neurons with significant differences in condition-dependent ΔF/F amplitude, the modified GLM analysis detects neurons based on condition-dependent response selectivity. To quantify this selectivity, we introduced a preference metric, the preference index (PI), which is proportional to the hyperbolic area of the ROC curve, calculated as (AUC - 0.5) × 2. The PI for neurons with temporal and directional selectivity was significantly higher than that for shuffled responses, particularly for delay-preferring neurons (Fig. 2C, D). Notably, more than 30% of the phase- and direction-selective neurons had a PI above 0.9, indicating a near-complete separation in their responses to the two visual stimuli (Fig. 2B-C). Since the PI was calculated based on averaged responses^19^, we also sought to quantify the consistency of the responses across trials. To this end, we constructed a reliability index (RI), which is proportional to the hyperbolic area of the auROC from the ΔF/F across all trials (see Methods). As with the PI, the RI was significantly higher in direction-selective neurons than in the shuffled control or non-directional neurons (Supplementary Fig. 3). Moreover, direction-selective delay neurons exhibited a greater RI than sample neurons, suggesting a refinement of the encoded information following stimulus offset (Supplementary Fig. 3D-I).

**Figure 2:**
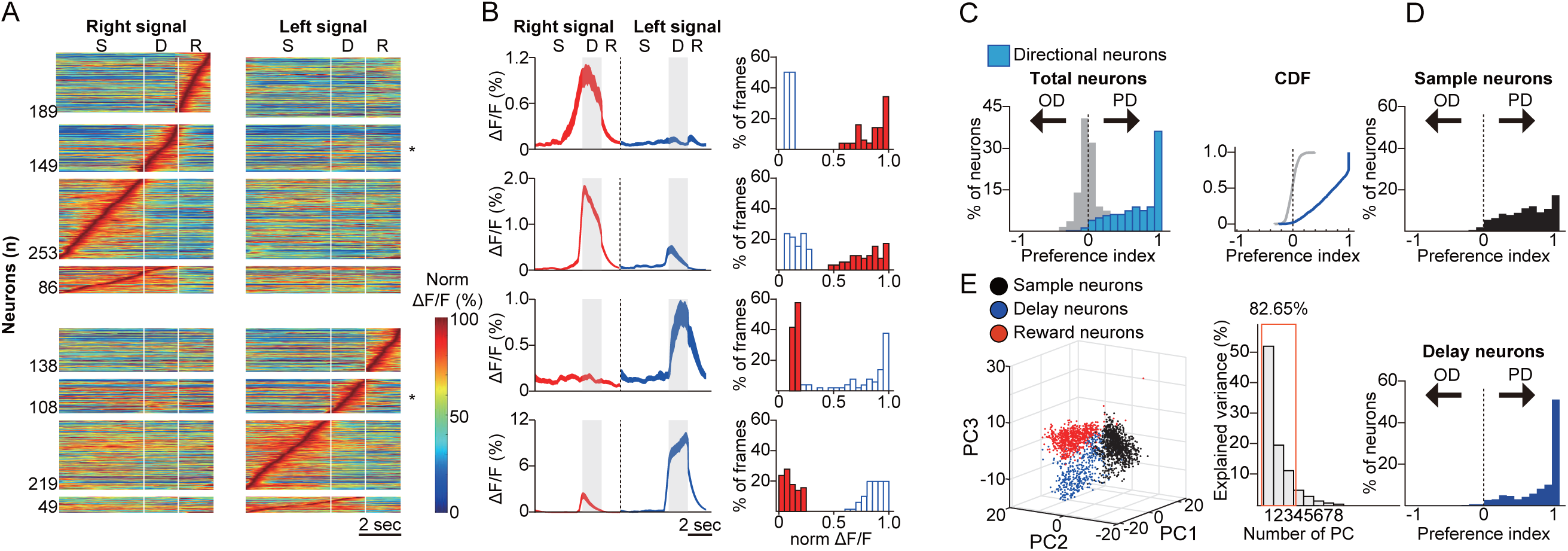
Identification and characterization of the direction- and phase-selective neurons. **A.** Color-coded trial-averaged ΔF/F of neurons with rightward (upper) or leftward (lower) directional phase selectivity. ΔF/F values were normalized to the maximum ΔF/F and aligned based on each neuron’s time-to-peak response. **B.** Example traces (left) and a histogram of trial-averaged ΔF/F from highly directional phase-selective neurons in the posterior parietal cortex (PPC). The thickness of the traces indicates the standard error. Blue and red in the histogram represent leftward and rightward trials, respectively. **C.** (Left) Histogram of preference indices (PI) for General Linear Model (GLM)-defined directional neurons (light blue) compared with those calculated from shuffled trials (gray). (Right) Cumulative distribution function (CDF) of PI values for the same data. **D.** PI histograms of directional neurons during the sample phase (upper) and delay phase (lower). **E** Principal components analysis (PCA) capturing the temporal activity patterns of individual neurons. GLM-defined selectivity for the sample (black), delay (blue), or reward (red) phase is indicated by colors. The variance explained by the principal components is also shown.

Phase-selective activities were further validated across the population using principal component analysis (PCA), which captured the temporal activity pattern of each neuron throughout the entire behavioral session (Fig. 2E). The GLM-identified phase-selective neurons (represented by dots with corresponding colors) form distinct populations within the principal component space, demonstrating that our modified GLM algorithm accurately identifies neurons with phase-specific firing patterns.

To determine whether the observed activity patterns were organized anatomically, we further assessed the relative co-localization of PPC neurons with similar response properties. Specifically, we compared the distances within groups of neurons with shared functional characteristics to those across groups (Supplementary Fig. 5). Consistent with previous findings from STM activity in navigation tasks^6^, we found no significant difference in the distances to functionally similar neurons compared to those with distinct firing patterns. This refutes the hypothesis that activity patterns are specifically localized. In summary, our results support the idea that activity sequences in the PPC can encode short-term memory of sensory input, even in a delayed match-to-sample task that does not require the encoding of heading direction or spatial location.

### Functional characteristics of directional neurons in error trials

Having established that PPC neurons exhibit direction- and phase-selectivity independent of spatial navigation, we next sought to investigate the mechanisms underlying the disruption of PPC activity during error trials. Specifically, we aimed to determine whether errors in STM could be attributed to inadequately encoded sensory inputs during the sample phase, poor retention of the encoded information during the delay phase, or a combination of both.

The activity of selective PPC neurons identified during correct trials exhibited markedly lower phase- or direction-selectivity during error trials (Fig. 3A). On an individual neuron level, we frequently observed that neurons with high PIs exhibited opposite directional selectivity during error trials (Fig. 3B). It is important to note that not all directionally selective neurons responded in the opposite direction, suggesting a mixed population response with some degree of mis-encoding in STM. There was a significant reduction in PI during incorrect trials, particularly among neurons that exhibited high PI during correct trials (Fig. 3C). In fact, the magnitude of PI reduction was greater for neurons that had high PI in correct trials (Fig. 3D and Supplementary Fig. 6). The median ΔPI for neurons with an initial PI between 0.9 and 1 was approximately -1, indicating that nearly 50% of the highly reliable neurons either lost response selectivity or responded to the opposite direction during error trials. To assess how these activity changes were related to the consistency of responses across individual trials, we examined the trial accuracy-dependent difference in RI values for phase- and direction-selective neurons as well as non-direction-selective neurons (Fig. 4A-I and Supplementary Fig. 6). This analysis revealed that a greater proportion of sample-phase neurons maintained directional information regardless of trial accuracy, while delay-phase neurons exhibited increased activation in the opposite direction during error trials. Notably, highly reliable directional delay neurons exhibited the greatest reductions in PI and RI during error trials (Fig. 4B-I). Together, these results suggest that highly reliable directional delay neurons are more likely to show a response pattern opposite to that observed during correct trials.

**Figure 3.**
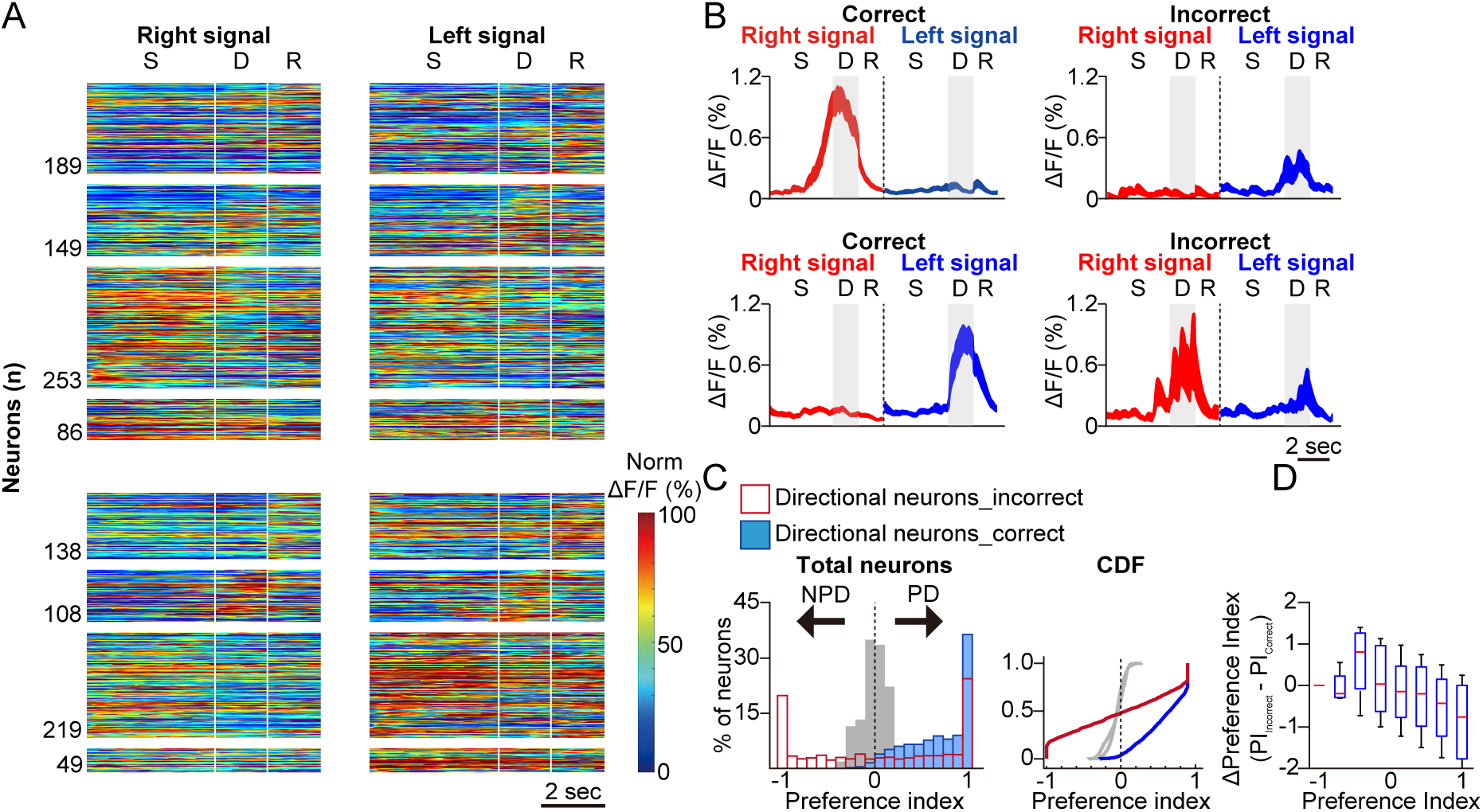
Responses of the direction- and phase-selective neurons in error trials. **A.** Same as Fig. 2A but in error trials. **B.** The same example neurons shown in the first and third row in Fig. 2B (left) and their mean traces in error trials. **C.** (Left) Histogram of preference indices (PI) of General Linear Model (GLM)-defined directional neurons in correct (light blue), error (red open), and shuffled (gray) trials. (Right) Cumulative distribution function (CDF) of the left histogram. **D.** A box plot illustrating the difference of PIs (ΔPI) between error and correct trials for each neuron, as a function of PIs in correct trials.

**Figure 4.**
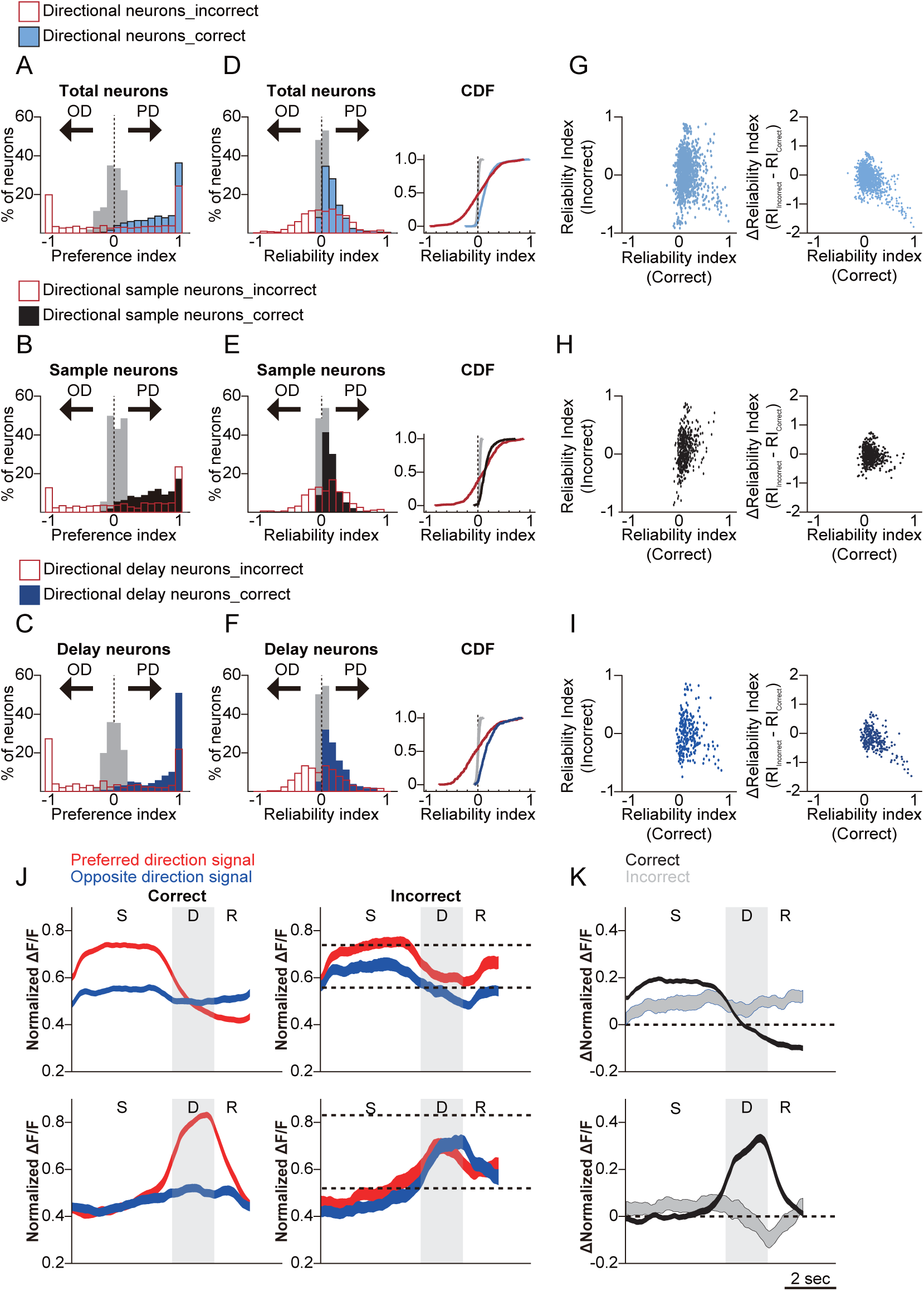
Comparison of responses between delay phase-selective neurons and sample phase-selective neurons during error trials. **A.** Histogram of preference indices (PI) for GLM-defined directional single-phase selective neurons during correct trials (light blue), error trials (red open), and shuffled trials (gray). **B.** Histogram of PI values for directional sample-phase selective neurons during correct trials (black), error trials (red open), and shuffled trials (gray). **C.** Histogram of PI values for directional delay-phase selective neurons during correct trials (blue), error trials (red open), and shuffled trials (gray). **D.** (Left) Histogram of reliability indices (RI) for GLM-defined directional single-phase selective neurons during correct trials (light blue), error trials (red open), and shuffled trials (gray). (Right) Cumulative distribution function (CDF) of the RI values from the left histogram. **E.** (Left) RI histogram for directional sample-phase selective neurons during correct trials (black), error trials (red open), and shuffled trials (gray). (Right) of the RI values from the left histogram. **F.** Left) RI histogram for directional delay-phase selective neurons during correct trials (blue), error trials (red open), and shuffled trials (gray). (Right) CDF of the RI values from the left histogram. **G.** (Left) Scatter plot comparing the reliability index RI in correct and incorrect trials for GLM-defined directional neurons in the posterior parietal cortex (PPC). RI values close to 1 during “correct” trials indicate significant responses in the preferred direction, while values close to -1 during “incorrect” trials suggest responses closer to the opposite direction. (Right) Scatter plot comparing the RI during correct trials with the difference in RI between incorrect and correct trials. ΔRI values closer to -1 indicate that responses during incorrect trials were closer to the opposite direction compared to correct trials. **H.** (Left) Scatter plot comparing the RI of GLM-defined directional sample-phase selective neurons in the PPC during correct and incorrect trials. (Right) Scatter plot comparing the RI (Correct) of GLM-defined direction-selective sample neurons with the difference between the reliability indices for incorrect and correct trials. **I.** (Left) Scatter plot comparing the RI of GLM-defined directional delay-selective neurons in the PPC during correct and incorrect trials. (Right) Scatter plot comparing the RI (Correct) of GLM-defined direction-selective delay neurons with the difference between the reliability indices for incorrect and correct trials. **J.** (Upper) Mean normalized ΔF/F of direction-selective sample-phase neurons in response to the preferred (red) and opposite (blue) directional signal during correct (left) and incorrect (right) trials. (Lower) Same as the upper plots but for direction-selective delay-phase neurons. **K.** (Upper) Δ mean normalized ΔF/F of direction-selective sample-phase neurons during correct trials (black) and incorrect trials (gray), calculated by subtracting the value of the opposite directional signal response from the preferred directional signal response. (Lower) (Upper) Δ mean normalized ΔF/F of direction-selective delay-phase neurons during correct trials (black) or incorrect trials (gray), calculated by subtracting the value of the opposite directional signal from the preferred directional signal.

A reversal in firing responses for neurons with high selectivity would predict a diminished overall difference between preferred and non-preferred directions. Indeed, we observed a significant reduction in the difference between ΔF/F of preferred and opposite directional cues during error trials (Fig. 4J). This reduced difference was the most pronounced among delay-selective neurons. Specifically, we noted a decrease in ΔF/F in the preferred direction and an increase in ΔF/F in the opposite direction, effectively rendering the two responses during the delay period indistinguishable (Fig. 4K). These findings suggest that the response selectivity of delay neurons was more profoundly affected than that of sample neurons during error trials.

### Activity trajectory distances of the direction-selective PPC neurons during the task

The observation that a subset of direction-selective neurons exhibited activity patterns corresponding to their response to the opposite directional signal during error trials suggests the possibility of either mis-encoding or disrupted retention of STM information. To directly compare the contributions of these two potential mechanisms, we next examined the temporal changes in the population-wide activity of directional neurons. Specifically, we aimed to determine whether the STM-associated population dynamics initially resembled the opposite direction during the sample phase, gradually shifted towards the opposite direction as the trial progressed, or reflected both patterns.

To this end, we first captured the dynamics of direction-selective neurons in a reduced dimension by representing their activity trajectories through PCA. The activity trajectories corresponding to correct preferred direction responses (specifically, the left trial responses from left-preferring neurons and the right trial responses from right-preferring neurons) were designated as preferred directional correct responses, or PP. In contrast, the trajectories corresponding to correct opposite-directional responses (i.e., right trial responses from left-preferring neurons and left trial responses from right-preferring neurons) were designated as opposite directional responses, or OO (Fig. 5 and Supplementary Fig. 8). We then calculated the distance between these trajectories as a metric of activity differences across conditions (Fig. 5A). The trajectory distance between PP and OO (d1) remained consistently high throughout the sample period and further increased during the delay period (Fig. 5C, d1). As a control, we also calculated the distance between randomly selected preferred trials and the remaining preferred trials to assess the magnitude of variability in activity (Fig. 5C, d0).

**Figure 5.**
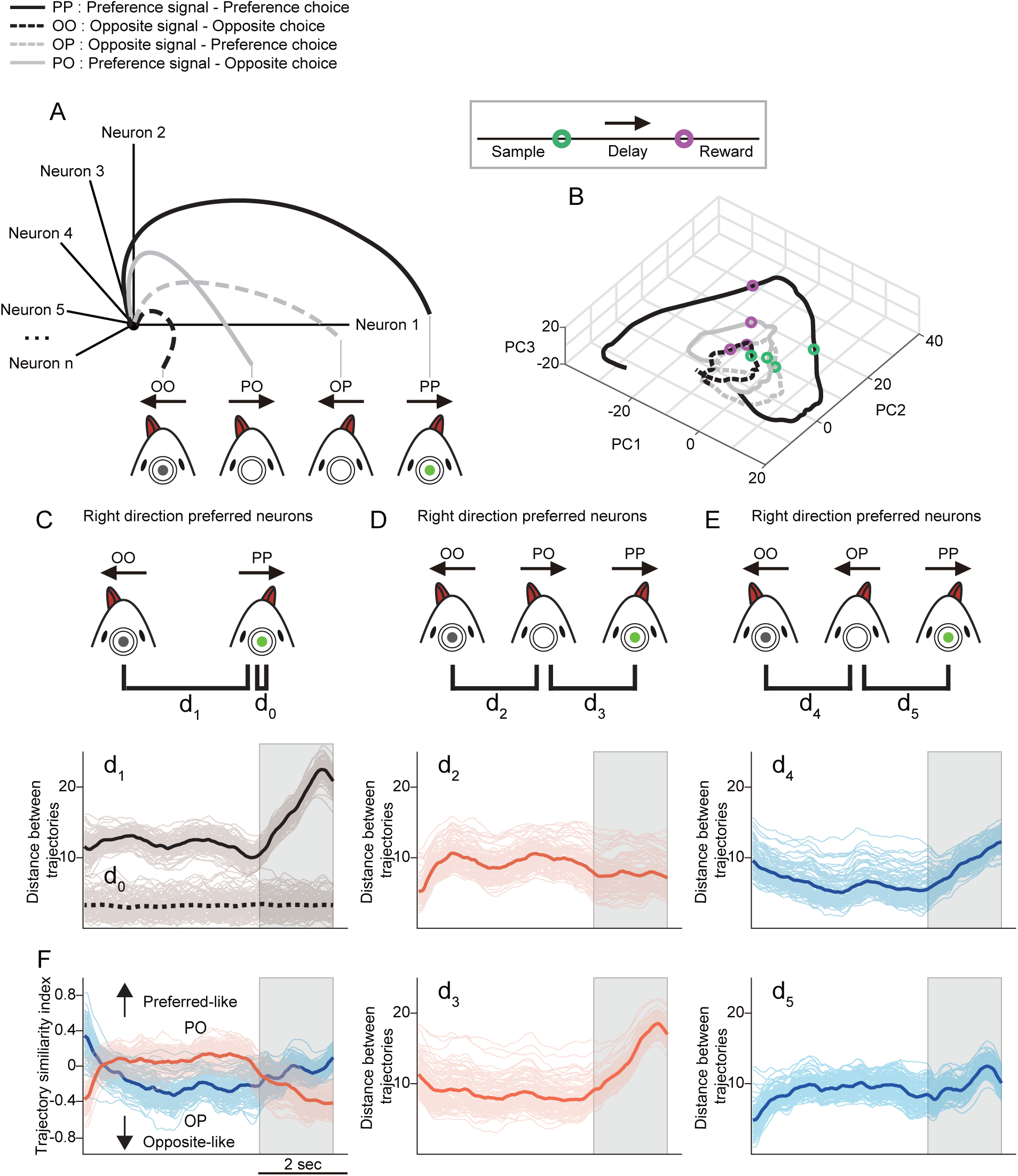
Differences in population activity of direction-selective neurons in correct and error trials. **A.** Schematic representation of the dimensionality reduction approach used to assess differences in population activity. **B.** Population activity trajectories of direction-selective neurons in response to preferred (dark colors) and opposite (light colors) directional stimulation during correct (blue) and incorrect (red) trials. The trajectories were visualized in a three-dimensional space for clarity, although analyses were conducted in a five-dimensional subspace. Eigenvectors were generated using 50% of the sorted neurons, and the remaining 50% were projected onto these eigenvectors. Faint lines represent values from 100 random iterations. **C.** (Upper) Schematic comparing trajectories of correct preferred direction responses (PP) and correct opposite direction responses (OO) for each neuron. (Lower) Temporal evolution of the Euclidean distance between PP and OO trajectories (d1). For comparison, d0 represents the trial-by-trial activity variation, measuring the distance between a randomly selected preferred direction response and other preferred direction responses within correct trials. **D.** d2: Euclidean distance between trajectories of correct opposite direction responses (OO) and incorrect preferred direction responses (PO). d3: Distance between trajectories of correct preferred direction responses (PP) and incorrect preferred direction responses (PO). **E.** d4: Distance between trajectories of correct opposite direction responses (OO) and incorrect opposite direction responses (OP). d5: Distance between trajectories of correct preferred direction responses (PP) and incorrect opposite direction responses (OP). **F.** Temporal dynamics of the trajectory similarity index for incorrect preferred responses (orange) and incorrect opposite responses (blue). The trajectory similarity index captures the degree of overlap between trajectories in the reduced-dimensional subspace (see Supplementary Fig. 8 for additional details).

To assess the changes in the activity of direction-selective neurons during error trials, we compared the trajectory distances from error trials to those from correct trials, PP and OO. In error trials where preferred directional inputs resulted in an opposite direction response (PO)— such as left-preferred neurons responding to left stimuli with a rightward response, or right-preferred neurons responding to right stimuli with a leftward response—the trajectory distance to the correct OO activity (Fig. 5D, d2) increased and saturated around the first 0.5 s of the sample period. However, the trajectory distance (d2) in the sample phase saturated at a lower value than d1, suggesting a relatively poor distinction in activity based on stimulus direction (two-tailed unpaired *t*-test; n=100, *p* < 0.001, Supplementary Fig. 8A). Subsequently, d2 decreased from the late sample phase into the delay period (gray area). In contrast, the trajectory distance between PO and PP (Fig. 5D, d3) decreased during the sample period, yet remained significantly greater than d0 (two-tailed unpaired *t*-test; n=100, *p* < 0.001, Supplementary Fig. 8B). This distance then increased dramatically during the delay phase. Taken together, these results suggest that during the sample phase of PO errors, PPC neurons fired similarly to PP neurons but with relatively poor distinction by stimulus direction. From the late sample phase to the delay phase, the activity increasingly resembled the opposite directional firing pattern. Similarly, in error trials where opposite stimuli resulted in preferred directional responses (OP), we observed that the activity showed decreasing distance to OO during the sample phase and drifted significantly during the delay phase (Fig. 5E, d4). The activity distance between OP and PP (d5) increased during the sample phase, followed by a slight decrease toward the end of the delay phase.

However, since the error activity trajectories may not necessarily lie within the same activity plane as the PP and OO trajectories, a closer distance to PP does not inherently imply a greater distance from OO. To assess the relative distances in a multi-dimensional space, we introduced a trajectory similarity index (TSI). The TSI quantifies the differences in distances to PP and OO, normalized by the sum of those distances, such that a higher absolute TSI indicates greater similarity. The polarity of the TSI value reflects whether the activity is more similar to PP (positive) or OO (negative) (illustration in Supplementary Fig. 8C; see Methods for details). Consistent with the analysis of distances to PP and OO activity (d2–d5), the TSI during the sample phase showed weak but significant directional activity in response to the given stimuli. As the delay phase progressed, however, the activity drifted toward the opposite direction (Fig. 5F).

### Population activity of PPC neurons during the task

The activity trajectories in the principal component dimension were computed using the combined activity of direction-selective neurons across sessions. To investigate how these dynamics relate to overall population activity, we applied measures that assess activity across all recorded neurons. Specifically, we used linear discriminant analysis (LDA) to compare population activity during error trials with that observed in correct trials, where the stimulus direction was either the same or opposite. This method provided a measure of the signal-to-noise ratio (SNR) of the encoded signal across the population, where the signal represents the distance between the two responses in the activity space, and the noise accounts for the variance within the responses (see Methods). We measured the SNR changes in error trials within a moving window of 50 frames in the hyperplane that best discriminates between correct left- and right-trials (Fig. 6). Consistent with the trajectory distances of direction-selective neurons, the SNR between correct and same-stimuli error trials (correct left trials [LL] vs. left stimulus-right response [LR], and correct right trials [RR] vs. right stimulus-left response [RL]) initially decreased, reaching saturation around the middle of the sample period (approximately 30 frames). As the frames from the delay period were included (after the 70th frame), the SNR increased (Fig. 6A, left). In contrast, the SNR between correct and incorrect trials with the opposite stimuli (same response error, LL vs. RL, and RR vs. LR) exhibited an opposite pattern: an initial increase and saturation over 30 frames during the sample period, followed by a subsequent decrease during the delay period (Fig. 6A, right). The patterns of same-stimuli and same-response errors aligned with the TSI changes in OP and PO errors, respectively (Fig. 5F). While grouping 50 frames across trials ensures separation between groups, this analysis obscures the SNR changes during the delay period. To better visualize the SNR changes, we replotted the measured SNR for every 30 frames within the delay period (Fig. 6B). While same-stimuli errors became more differentiated during the delay phase, a decreasing SNR was observed in same-response errors. Across all frames, during same-stimuli errors, PPC activity was relatively similar in the sample phase but diverged more significantly during the delay phase. In contrast, in same-response errors, the activity differences driven by stimulus direction in the sample phase diminished during the delay phase (Fig. 6C). These results underscore the role of activity drift toward the opposite direction in contributing to erroneous STM.

**Figure 6.**
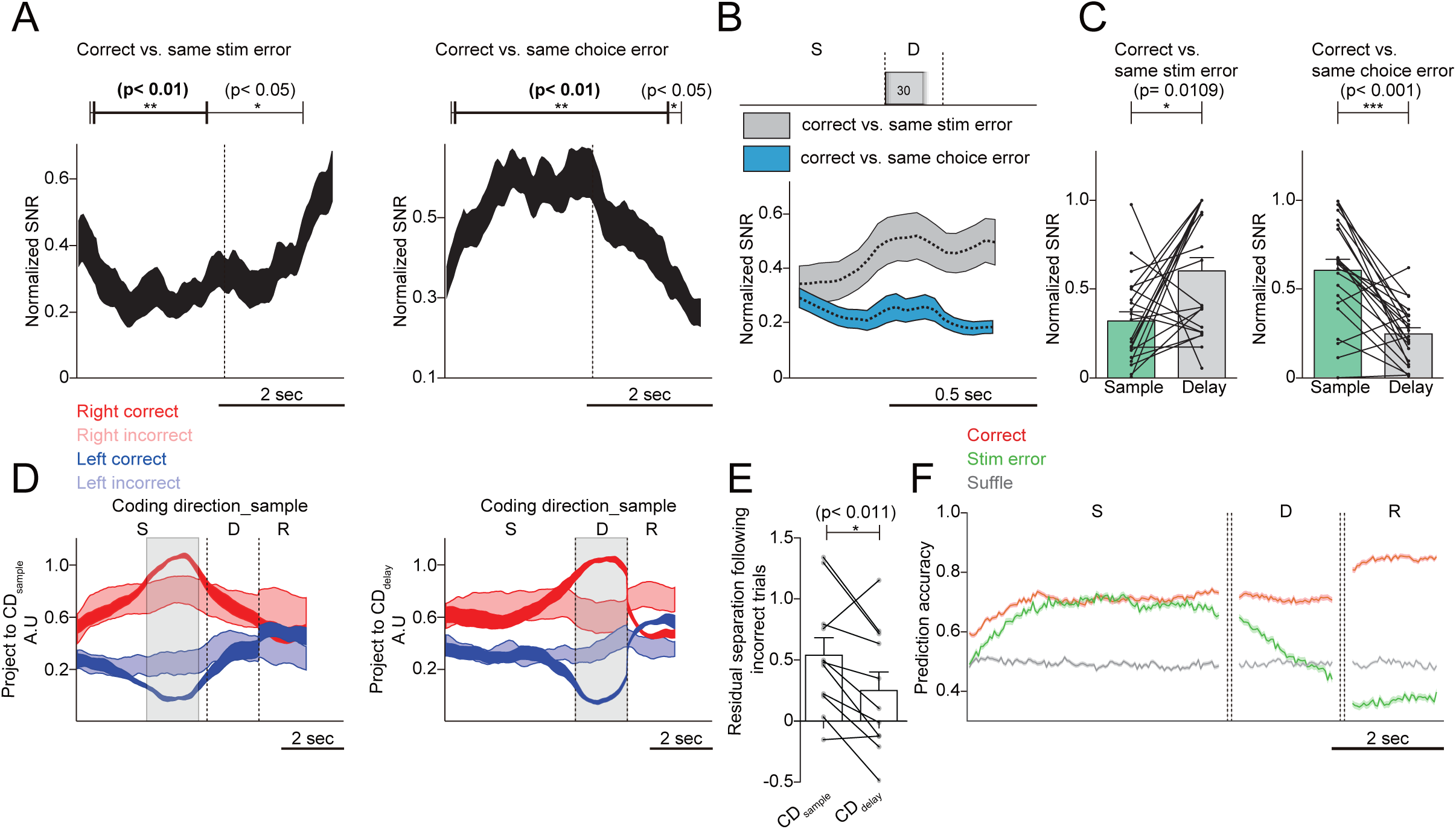
Population dynamics of posterior parietal cortex (PPC) neurons in correct and incorrect trials. **A.** Time-dependent changes in the normalized signal-to-noise ratio (SNR) to distinguish neural activity responses in correct and error trials using linear discriminant analysis (LDA). SNR was calculated using a sliding window of 50 frames, advancing one frame at a time. The vertical dashed line marks the initiation of the delay phase. **Left Panel:** Mean normalized SNR between neural activity clusters for correct trials and same-stimulation error trials (e.g., LL vs. LR and RR vs. RL) as a function of time. Paired *t*-tests revealed significant differences: black horizontal line (frames 8–107, *p* < 0.05) and bold black line (frames 11–65, *p* < 0.01). **Right Panel:** Mean normalized SNR between correct trials and same-choice error trials (e.g., LL vs. RL and RR vs. LR). Significant differences are indicated by black line (frames 3–112, *p* < 0.05) and bold black line (frames 6–105, *p* < 0.01). **Key:** Line thickness indicates mean ± SEM across sessions. **B.** Mean normalized SNR of LDA results during the delay phase. **Gray Line:** Results comparing correct trials and same-stimulation error trials. **Blue Line:** Results comparing correct trials and same-choice error trials. Line thickness represents the standard error across sessions. **C.** Comparison of normalized SNR values during the sample phase (frames 66–115) and the delay phase (frames 121–170). **Left Panel:** Correct vs. same-stimulation error trials. **Right Panel:** Correct vs. same-choice error trials. Statistical significance was determined using paired *t*-tests, denoted as *p* < 0.05*, p* < 0.01, and *p* < 0.001. Error bars indicate mean ± SEM across sessions. **D.** Projections of population activity along two coding dimensions: **CDsample** (coding dimension for the sample phase) and **CDdelay** (coding dimension for the delay phase). Projections are shown for each direction (red: right; blue: left) and response accuracy (dark: correct; light: incorrect). Line thickness represents mean ± SEM across sessions. **E.** Residual separation defined as (Right incorrect - Left incorrect) / (Right correct - Left correct). (paired *t*-test, *p* < 0.05). **F.** Accuracy of neural activity prediction over time, decoded using a support vector machine (SVM). Line thickness indicates mean ± SEM across sessions.

While our analysis using PCA and LDA revealed activity differences between error and correct trials, the observed activity differences may not directly reflect the representation of the directional content of sensory inputs. To address this, we employed a dimensionality reduction approach to identify population activity modes that distinguish relevant features, particularly the signal corresponding to the direction of motion in the visual stimulus (coding direction or CD). To directly compare signal direction-dependent activity modes, we computed projections along a vector in the activity space of all recorded PPC neurons that maximally differentiate sensory direction in the sample phase (CDsample) and the delay phase (CDdelay) (Fig. 6D). To compare the separation of directional signals, we projected the error trial activities onto this vector. In comparison to correct trials, the directional sensory signals were weaker and less distinguishable between directions, particularly during the delay phase of error trials (Fig. 6D-E).

Consequently, we aimed to assess how accurately correct or error trials could be predicted. We trained a Support Vector Machine (SVM) model using the activities of PPC neurons during correct trials. Subsequently, we evaluated the accuracy of decoding sensory direction from the trial-by-trial population activity of PPC neurons using the SVM decoder. PPC activity from randomly selected 70% of correct trials in a session was labeled as belonging to either the left or right direction, and the remaining trials were used to test prediction accuracy (Fig. 6F). For correct trials, the prediction accuracy of the classifier significantly increased above the shuffled control level (gray line) at the onset of the sample phase and remained elevated until the end of the delay period. When assessing error trials using the same hyperplane generated from correct trials, the prediction accuracy showed a slight temporal delay but eventually reached a level comparable to correct trials. This suggests that despite a significant reduction in the distinction between stimulus directions, the population activity of PPC neurons still encodes the correct direction. However, during the delay period of error trials, the prediction accuracy dropped rapidly, falling below the chance level by the end, indicating a tendency toward choosing the opposite direction. Taken together, our results demonstrate that population activity in the PPC during error trials, especially in the delay phase, resembles the responses to the opposite directional signal, suggesting that STM memory errors are associated with a reversal in stimulus content encoding within the PPC.

## Discussion

### Distinguished sensory information maintenance from navigational and movement signals in PPC activity during STM

In the current study, we observed propagating neural activity that tracked the sensory information being maintained during STM task performance. Similar observations have previously been reported in STM-based virtual T-maze tasks^6^. However, ^6^subsequent studies have questioned the interpretation of these findings, suggesting that choice-dependent activity sequences could be predicted by heading direction and the animal’s position in the virtual corridor, potentially reflecting a navigational signal instead of STM processing^6,13^. This perspective aligns with earlier studies demonstrating that PPC neurons exhibit activity changes associated with locomotor actions, such as turning and forward motion, particularly when these movements are linked to spatial references^11,12,20,21^. Distinguishing between PPC activity related to sensory information maintenance and that influenced by navigational context is therefore critical for a more accurate understanding of PPC function in STM. Sensory cue-dependent go/no-go tasks also present interpretational challenges, as the observed activity may reflect preparatory signals related to impulsive licking behavior or its suppression, rather than sensory maintenance^6,14,19,22^. In our study, we re-examined PPC activity during an STM task that required retaining visual information and making directional choices based on equivalent licking behavior, but without incorporating a spatial component. Remarkably, we found similar activity patterns in PPC neurons, with direction-selective activity sequences that predicted sensory direction. These patterns were observed not only in the absence of navigational signals but also during the delay period when no ongoing stimulus was present. Furthermore, disruptions in PPC activity patterns during error trials were consistent with a drift toward an erroneous representation of the stimulus direction, further supporting the interpretation that PPC activity reflects an STM signal.

### Difference in population activity in error trials

A substantial proportion of PPC neurons exhibited significant directional preferences, with over 30% demonstrating exceptionally high selectivity (PI > 0.9, Fig. 2A-C). Most direction-selective neurons were tuned to specific phases, rather than maintaining sustained activity across both the sample and delay periods (Fig. 2A, D). This high directional selectivity, or high PI, may be facilitated by the wiring motifs in the PPC, where co-selective neurons form strong and more frequent synaptic connections while inhibiting anti-selective neurons^23^. In the PCA-driven activity subspace, which captured over 70% of the activity of selective neurons (Supplementary Fig. 8), trajectories corresponding to two directional visual stimuli began diverging immediately after stimulus onset and continued to separate throughout the delay period (Fig. 5C, 6B). Surprisingly, this differentiation persisted and even increased during the delay period, despite the absence of ongoing sensory stimuli. This observation aligns with the significantly greater direction selectivity observed in delay neurons compared to sample neurons (Fig. 2D). These findings support the hypothesis that the PPC primarily functions to maintain differentiated sensory inputs acquired during the sample period, rather than to detect and encode these signals. The latter function may instead rely on early sensory cortical processing ^6,14,24^. The progressive increase in activity divergence during the delay phase may reflect processes aimed at optimizing visual information retention while minimizing activity variability. This could occur concurrently with premotor neuron preparation for invariant motor responses ^25,26^. Further studies systematically comparing PPC activity under varying stimulus durations and complexities could clarify these possibilities.

Our complementary multivariate population analyses consistently revealed sustained population-level differences in activity in response to distinct stimuli during correct trials (Fig. 5C, 6F). Although this encoding pattern involves distributed activity across multiple neurons, it resembles the increased or decreased sustained firing rates observed in PPC neurons of primates^8,27^ and rats performing STM tasks^9^. These findings suggest that population activity differences in the mouse PPC may represent a form of evidence quality. Differences in PPC activity between cue directions that emerge during the sample period appear to underpin stimulus encoding, while their maintenance during the delay period reflects STM retention. Consistent with this model, we observed weaker directional differentiation and poorer maintenance of population activity differences during error trials. These results suggest that STM errors are associated with suboptimal sensory encoding and a gradual weakening of this information throughout the task (Fig. 5-6). However, despite reduced activity distinction and some neurons "misinterpreting" stimuli during the sample phase of error trials, population-level PPC activity still encoded the correct direction to a substantial extent (Fig. 6F).

### Increasing opposite directional activity of PPC neurons in error trials

The origin of the reduced distance in activity space observed during error trials remains unclear. However, changes in the animal’s arousal state may contribute to this diminished encoding. Arousal levels vary continuously, and neuronal responses in sensory cortices, as well as higher-order cortical areas such as the PPC, are critically modulated by attentional states^6, 6,28–33^. During error trials, insufficient arousal could lead to reduced PPC neuron activity in response to preferred visual stimuli, resulting in weaker separation in activity space (Fig. 3D, 4G-H).

Nevertheless, this explanation does not account for neurons with negative PI values—those that exhibit stronger firing to the opposite directional stimuli compared to the preferred direction. For these neurons, we propose that PPC activity encodes how the stimulus was ‘interpreted’ rather than its identity, particularly for delay neurons and, to a lesser extent, for directional sample neurons (Fig. 3, 4). This interpretation aligns with findings from macaque studies, where PPC neurons showed firing changes predictive of behavioral choices even when the stimulus contained 0% coherence in a random dot motion task ^8^.

Recent anatomical analyses have revealed that the probability and strength of connections among L2/3 PPC neurons depend on the similarity of their directional selectivity^23^. Furthermore^23^, PPC subnetworks with opposing selectivity appear to suppress each other through mutual inhibitory connections^23,23,34,35^. Building on these findings and our results, we propose that direction-selective delay neurons form stronger subnetworks than sample neurons. In error trials, although the opposite-directional PPC activity during the sample phase does not initially alter the perceived direction of the stimuli, this activity may be amplified through the tightly interconnected delay neurons. This amplification could eventually result in an inversion of the encoded direction, leading to an error. This hypothesis is supported by our findings, which show that delay neurons exhibit significantly higher PI and RI values compared to sample neurons (Fig. 2D, 4). Additionally, we observed a drift in population activity toward the opposite direction as the delay period progresses (Fig. 5F, 6).

### Limitations of the study

Due to the limited number of task-relevant neurons recorded in a single trial and the inherent noise in cortical activity, much of our analysis was based on averaged activity across numerous trials of the STM task. Consequently, it was not feasible to reliably track the onset and amplification of inappropriate directional firing in this study. As such, both the origin of erroneous activity within the subnetwork and the mechanisms underlying its amplification remain unresolved. Additionally, although PPC activity was recorded while mice were head-fixed to perform the task, unintended movements or fluctuations in arousal state may have occurred, potentially modulating PPC activity^33,36^. Future studies could explore^33,36^ trial-to-trial variability under varying attentional conditions using high-density electrophysiological recordings, which may provide deeper insights into these dynamics^37–41^.

### Conclusion

In conclusion, our study advances the understanding of the neural mechanisms underlying sensory encoding and retention in STM tasks, highlighting the pivotal role of the PPC in these processes. By elucidating the relationship between PPC activity patterns and task performance, we provide valuable insights into the cognitive functions of the PPC, with significant implications for understanding STM errors. Given that impaired STM performance is a hallmark symptom of schizophrenia^42–44^ and an endophenotype of other neuropsychiatric disorders^45^, understanding ^45^the nature of STM errors holds considerable translational relevance. Our findings pave the way for future research aimed at developing novel therapeutic approaches for these conditions. Moreover, exploring trial variability and attentional conditions with advanced recording techniques will deepen our understanding of PPC function in STM and its disruption in disease.

## Methods

### Animals

All animal experiments were approved by the Animal Experiment Ethics Committee of the Korea Brain Research Institute (KBRI) (approval no. IACUC-17-00031). The experiments were performed using male C57BL/6J-Tg(Thy1-GCaMP6f)GP5.5Dkim/J transgenic mice (n = 6).

### Cranial window and head plate preparation

Mice aged 8 to 12 weeks underwent cranial window and head plate implantation surgery. To alleviate pain and reduce inflammation, they received subcutaneous injections of ketoprofen (5 mg/kg) prior to surgery. Deep anesthesia was induced using a mixture of 1–2% isoflurane or a ketamine/xylazine solution (100 mg/kg and 10 mg/kg, respectively). Anesthesia was confirmed by the absence of the pedal withdrawal reflex. Both eyes were covered with petroleum jelly, and the incision site was shaved. The scalp was sterilized with ethanol and betadine before being excised. Using a scalpel, the periosteum was removed, and the skull surface was roughened with a drill burr (Miniature Carbide Burr Drill Bits Set, Stoelting) to ensure proper adhesion of the dental cement. The skull was rinsed with Ringer’s solution, and a craniotomy approximately 3 mm in diameter was performed over the posterior parietal cortex of the right hemisphere. The craniotomy was sealed with a double-layered glass coverslip (3/5 mm, Harvard Apparatus) combined with NOA71 adhesive (Norland) and secured with surgical adhesive (Vetbond, 3M). The dura mater remained intact. A tailor-made, commercially pure titanium head plate (rectangular, 19 mm x 12 mm x 1 mm) with a 4.5 mm central opening was affixed on top of the window using dental cement (Super-Bond C&B kits, Sun Medical, darkened with black pigment). After surgery, mice were placed in a warmed chamber for at least four hours for recovery and post-operative monitoring. Subsequently, the animals were housed individually in cages.

### Behavior task

After full recovery from surgery, the mice were provided with limited water (1 mL/day), and their body weights were monitored regularly. If a mouse’s body weight dropped below 75% of its initial weight, water restriction and training were halted. The mice were trained to perform a delayed match-to-sample task while their heads were fixed in place using a holder (modified MAG-1, Narishige Scientific Instrument, Tokyo, Japan). The visual signal for discrimination consisted of high-contrast sine-squared wave gratings moving either left/right or down/right, with specific spatial (3 cm/cycle) and temporal (2 Hz) frequencies. This signal was displayed on a monitor positioned directly in front of the mice at a distance of 25 cm. Custom-written software running on a Raspberry Pi 3 (RPi3) controlled both the stimuli generation and behavioral training. Mice were trained separately in either the left/right or down/right directions, and the data from both sets of trials were pooled and analyzed together.

The position of the drinking spouts was individually adjusted for each mouse. The lick detection circuit and reward solenoid were controlled by an Arduino Mega2560 (rev 3, Arduino), which was connected to the Raspberry Pi 3 (RPi3). During training, the mice earned the majority of their daily water intake (1 mL) through task performance. Each visual stimulus lasted for 4 s, followed by a 2-s pause. After this pause, the spouts quickly moved to a position accessible by the mouse’s tongue for 1 s. A successful lick (i.e., licking the left-hand spout in response to a leftward or downward-moving signal, or licking the right-hand spout in response to a rightward-moving signal) during this window resulted in a reward of approximately 5 µL of tap water. Incorrect licks were not rewarded and were followed by a 2-s time-out, after which the trial continued. After the reward period or an incorrect lick, the spout retracted quickly and stayed out of reach until the next trial, which began 4 or more seconds later. Training progressed in stages: Initially, the mice learned to lick a non-moving central lick port in response to either visual cue. Next, they were trained to match the visual direction with the corresponding reward spout. Subsequently, the delay was gradually increased by introducing moving lick ports. Once the mice consistently performed with over 70% accuracy at a 2-s delay, both behavioral performance and PPC activity were recorded for further analysis.

### *In vivo* two-photon calcium imaging

We performed two-photon imaging using a HyperScope (Scientifica) and ScanImage (MBF Bioscience) setup, powered by a Ti:Sapphire laser (Chameleon Ultra II, Coherent). The imaging system was configured to target the GCaMP6f calcium indicator, which was excited at 910 nm using a 16x water immersion objective lens (N16XLWD-PF, Nikon). A custom-made light barrier was attached to the objective lens to isolate the imaging system from any visual stimuli. Images were captured at a resolution of 512 × 512 pixels (700 µm × 700 µm region of interest [ROI]) at a frame rate of 30 Hz in layer 2/3 of the PPC. The average laser power at the objective lens was kept below 100 mW to minimize tissue damage. The imaging and behavioral systems were synchronized via a frame trigger signal output, which was counted by an Arduino Mega 2560 (rev 3).

### Image preprocessing and cell selection

Calcium imaging data were processed using the CaImAn software framework. Motion correction was applied in segments of 128 × 128 pixels, with each segment overlapping by 48 pixels to ensure continuity across frames. For neuronal source identification, the ‘greedy_roi’ method was employed, with expected neuron sizes set to 4 × 4 pixels for detailed imaging and 3 × 3 pixels for full-brain imaging. Only components with a signal-to-noise ratio (SNR) of at least 2.0 and 90% pixel correlation were considered reliable. Fluorescence signal decay rates were normalized by applying z-scoring. A decay rate of 3.0 s was used for indicators localized in the nucleus, and 1.0 s for those in the cytosol. To align neural activity with behavioral actions, all time series data were synchronized to a uniform sampling rate of 30 Hz, corresponding to the frame capture times of the camera and microscope.

### Automated event-related neuron detection in calcium imaging data

We used a generalized linear model (GLM) to regress recorded calcium signals against a time series of task events. We propose a novel method for automatically searching event-related neurons in calcium imaging data (Supplementary Fig. 9). Our method searches the ROI pixelwise using event timing information. By utilizing a statistical method to determine the relationship between the calcium trace of each pixel and the event timing, the searched ROI pixels are clustered. This process enables the automatic detection of event-related ROIs without requiring any additional detection procedures while also enabling the detection of components with low SNR or fine size as ROI.

The proposed method determines the event relevance of calcium traces to detect event-related neurons in calcium imaging data. Rather than relying on the morphological shape of the neuron for ROI detection, the proposed method determines the event relevance of each pixel’s calcium trace independently. If ROI detection is performed by reflecting the neuron’s morphological shape, the detection performance may decrease with low SNR or fine-size components. The proposed method utilizes a generalized linear model (GLM) to detect ROIs by assessing the relationship between the calcium trace of each pixel and the calcium signal model that is strongly associated with the event. To create a calcium signal model that is strongly associated with events, a temporal component model of calcium dynamics and event timing is necessary. Event timing refers to the points in time when tasks or stimuli occur, and the calcium signal of event-related neurons should show a clear response at these times. In contrast, non-event timing refers to periods of spontaneous activity that are unrelated to the event. Neurons associated with events exhibit strong activation at event timing and minimal activation at non-event times. Therefore, we define a spike train 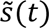 of the calcium signal most strongly associated with the event:

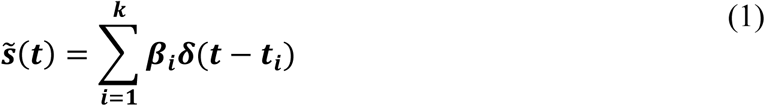

where 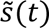 is composed of the product of the *β* (amplitude of the spike) and the impulse train. An event-related neuron is expected to activate at least once after an event. Denoting the *i^th^* event timing by 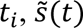 becomes a spike train of event-related neurons with an amplitude of *β*_*i*_ at event timings.

The temporal component of calcium dynamics is determined by the activity of a calcium indicator. When the calcium indicator is combined with calcium, luminous intensity changes. The fluorescence trace of the calcium indicator was modeled in the form of a double exponential and is expressed as:

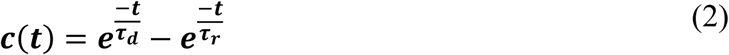

where *τ*_*r*_ and *τ*_*d*_ are the time constants for the increase and decrease of fluorescence, respectively^46^.

We express the modeled event-related calcium trace 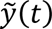 of the neuron by the fluorescence trace model of the calcium indicator *c*(*t*) with the convolution of the spike train 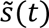.

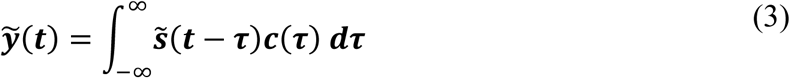

We defined the event-related calcium trace model 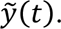 The ROI was detected by determining the relationship between this model and the pixel’s calcium trace *y*(*t*) using the least square method and 1-sampled *t*-test. Through the least square method, it is possible to find the estimator of the regression coefficient *β* (the amplitude of the spike in equation (1)) that minimizes the residual of the pixel’s calcium trace *y*(*t*) and the event-related calcium trace model 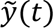 of the neuron. The *β* estimator matrix 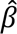 up to the kth event was expressed as:

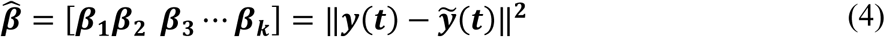

Notice that 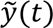 has a weight *c*(*t*) for event timing. By *c*(*t*), as the event timing and activation timing are closer, the value of the regression coefficient *β* increases. It was determined whether a neuron is event-related through the distribution of *β*. As the mean of 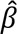 is close to 0 or the variance of 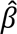 is larger, the activation timing of neurons has little relation to the event timing. Therefore, we tested the null hypothesis that β is extracted from a normal distribution with a zero mean and unknown variance, using a 1-sampled *t*-test. At this time, the test statistic is as shown:

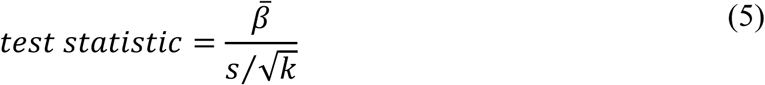

where 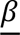 is the mean of the 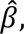 *s* is the standard deviation of the 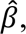 and *k* is the size of 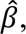 also known as the trial number of events. The proposed method uses a *p*-value of 0.05 to determine a pixel with a statistically significant difference from a pixel of an event-related neuron. When the event association determination is completed for all pixels of the calcium image, the proposed method finally defines the event-related ROI by clustering the pixels associated with the event. The proposed method clusters pixels to ROIs automatically, where the clustering is performed based on the Moore-Neighbor tracking algorithm and determines connectivity between pixels ^47^. Finally, the proposed method provides ROI clusters corresponding to event-related neurons (Supplementary Fig. 9).

### Selectivity

To identify neurons activated during specific phases of different behavioral patterns, we employed a uniquely developed GLM. To validate these findings, we introduced a preference index (PI) calculated via a receiver operating characteristic (ROC) analysis. This analysis assesses the theoretical observer’s ability to distinguish between different trial types based on neural responses. The preference index is determined from the area under the ROC curve (auROC) using the formula: 2 × (*auROC* − 0.5), producing a range from -1 to 1. We applied this index to segments representing each behavioral phase, using average neural activity across trials to compute specific values. The preference index (PI) compares trial-averaged neural activity between right and left conditions. In addition, the reliability index (RI) is calculated by comparing neural activity across all individual trials between the two conditions, without averaging, to assess the consistency of the neuron’s response. Lastly, the phase selective index (PSI) is derived by comparing the trial-averaged neural activity during the phase of interest with that from non-related phases. By using these indices, we were able to quantitatively assess the degree to which individual neurons preferentially respond to different trial types, offering insights into their roles within the observed behavioral patterns.

### PCA

We used Principal Component Analysis (PCA) on neurons sorted using the GLM. The goal was to infer an intrinsic manifold, representing the collective neuronal activity. We calculated the variance explained by the principal components of correct trials, observing that five principal components accounted for a similar proportion of variance in right-preferring, left-preferring, and all directional neurons (right-preferring neurons: 72.4%, left-preferring neurons: 69.0%, all directional neurons: 70.4%). The PCA procedure for analyzing PPC neuron data followed these steps: We structured the optical data from each experiment into an *X* = *M* × *N* matrix, where *M* is the number of frames per trial, and *N* is the number of sorted neurons in a session (*N* = *N*_*sample*_ + *N*_*delay*_ + *N*_*reward*_). Directional neurons (both right-preferring and left-preferring) were grouped together for subsequent analysis. Prior to conducting PCA, we aligned the neurons according to their preferred direction. To ensure comparability across neurons, we applied z-score normalization to each column of the data matrix. This process adjusts the data to have a mean of 0 and a standard deviation of 1, making the PCA more robust to scale variations between neurons.

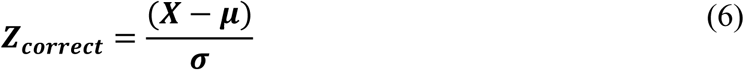

Where *μ* is the mean and *σ* is the standard deviation of neuron activity.

We calculated the covariance matrix, *C*, from the normalization data *Z*, to understand how the variables relate to one another.

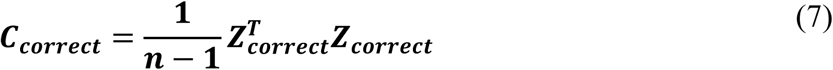

Where *n* is the number of neurons.

To further analyze the neuronal activity data, we computed the eigenvalues and eigenvectors of the covariance matrix *C*. The eigenvectors correspond to the directions of maximum variance in the data, while the eigenvalues indicate the magnitude of these variances. Neuronal activity data were projected onto the principal components to transform the data into a new space defined by these components. To compute relative changes in incorrect trials compared to correct trials, we projected the incorrect trials onto the eigenvector induced by correct trials and quantified the variations through the distances between trajectories.

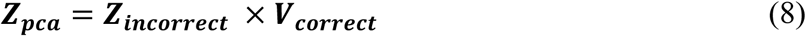

Where *V*_*correct*_ represents eigenvectors and *Z*_*pca*_ represents the data transformed into the principal component space.

### Coding direction

This procedure closely follows previously described methodologies ^48,49^. The activity of *n* neurons, recorded simultaneously during the entire session, was analyzed (Fig. 6). The population activity of *n* neurons formed a trajectory in the *n* - dimensional activity space during each session, with each dimension representing the activity of a single neuron. In this activity space, we identified directions (or modes) that maximally separated the neuronal pathways associated with different trial conditions and specific phases. For trials with lick-correct left and lick-correct right instructions, the trial-averaged activity differences of the *n* neurons were computed within a 1.7-s window (frames 121–170) following the visual cue offset. This resulted in the delay mode (*CD*_*delay*_), which was calculated as follows:

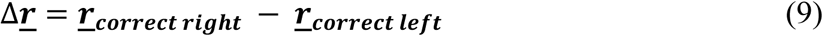

The resulting vector was normalized by its *l*^2^ norm, yielding a weight vector, with one weight assigned to each neuron *i*, as follows:

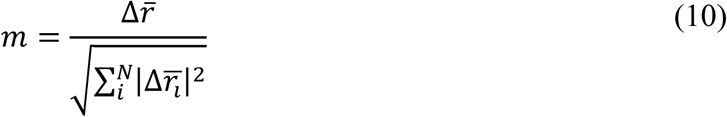

Projections of the neural activity onto the mode over time were computed as follows:

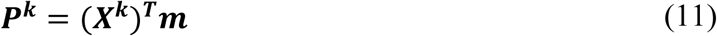

Where *X*^*k*^ is the *N* × *T* matrix of averaged activity of neurons over time for session *k*.

To maintain consistency in the analysis despite the varying number of neurons recorded in each trial, we normalized each mode by its *l*^2^ norm, as indicated in equation (10). This normalization ensured that the projected neural activity, *p*^*k*^ remained unaffected by the quantity of neurons recorded simultaneously. In instances where a specific neuron did not show any specific activity during a particular trial, we assigned a weight of zero to that neuron in both equations (10) and (11), for calculating the projected activity for that trial. This approach guaranteed that the analysis could accurately reflect the contribution of active neurons in each trial, without skewing results due to differences in neuron sampling across trials.

The sample mode (*CD*_*sample*_) and the reward mode (*CD*_*reward*_) were defined to capture distinct aspects of neural activity corresponding to different phases of the trials, specifically focusing on the behavioral outcomes of licking correctly to the right or left.

*CD*_*sample*_: This mode was determined by constructing an *n* × 1 vector representing the differences in the average activity of *n* neurons during trials that resulted in correct licks to the right or left. These differences were averaged over a 1.7-s window (frames 56–115) coinciding with the beginning of the stimulus presentation. By focusing on this early window, *CD*_*sample*_ captures the neural activity patterns triggered by the initial stimulus, reflecting how neurons differentiate between trial types in response to the stimulus.

*CD*_*reward*_: Similarly, this mode was defined by an *n* × 1 vector, but it focused on the average activity differences of *n* neurons during trials with correct licks to the right or left, within a 1-s window (from frame 171 to frame 200) at the conclusion of the delay phase. The *CD*_*reward*_ specifically targets the neural response patterns associated with the anticipation or receipt of a reward, highlighting how neuronal activity aligns with behavioral outcomes following the stimulus-response interval.

In addition to the selective modes that focused on differentiating neural activity based on specific trial outcomes, we identified a nonselective ramping mode (*D*_*reward*_). This mode was designed to capture general changes in neural activity towards the end of the delay phase, irrespective of the trial outcome (whether the lick was correct to the right or left). To compute this ramping mode, we adopted a methodology similar to that used for the reward mode. However, instead of distinguishing between trial outcomes, we aggregated neural responses from both correct-right and correct-left licks. We then calculated the difference in trial-averaged activity during the last second of the delay phase (from frame 141 to frame 170) compared to the first second of the reward phase (frames 171–200).

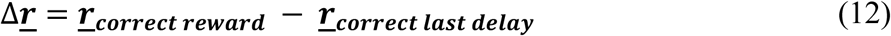

To facilitate a direct comparison between the sample and delay phases while considering their distinct neural dynamics, we employed the Gram-Schmidt process for orthogonalization of these phases in a specific sequence^50^. This process was applied to the modes in two different orders: first, starting with the *CD*_*reward*_, then the *D*_*reward*_, followed by the *CD*_*sample*_; and second, beginning with the *CD*_*reward*_, then the *D*_*reward*_, and finally, the *CD*_*delay*_. Orthogonalization using the Gram-Schmidt process ensures that each mode represents a unique aspect of neural activity, independent of the others, thus allowing for a clearer analysis of the distinct contributions of sample and delay phases to overall neural dynamics.

Furthermore, to assess the neural activity’s alignment with behavioral outcomes - specifically, the differences between correct and incorrect responses - we projected the trial-averaged activity from incorrect trials onto the modes defined by correct trials across sessions. This projection allowed us to compare the neural activity associated with correct and incorrect trials within the same mode, providing insights into how specific patterns of neural dynamics correlate with behavioral performance. This approach helped in understanding the neural underpinnings of decision-making and error processing by quantifying the extent to which neural activity during incorrect trials deviates from the established patterns of correct trials.

After projecting the neural activity data along the direction defined by CD (which represents the difference in neural activity between conditions), we normalized the resulting projections. This normalization involved adjusting the projections based on the average value obtained from correct rightward trials during the specific phase associated with each mode, for every session. This step ensured that the comparison across different trials and sessions was on a consistent scale, facilitating a more accurate analysis of the neural activity patterns.

To further investigate the differences in neural activity patterns between incorrect and correct trials, we introduced the concept of "residual separation." This metric quantifies the deviation in the neural trajectories of incorrect trials from those of correct trials. The residual separation effectively captures the extent to which the neural representation of incorrect trials diverges from the expected pattern observed in correct trials, providing insights into the neural mechanisms underlying error processing and decision-making accuracy. This analysis allowed us to better understand how the brain’s neural activity aligns with behavioral outcomes and how it adapts or fails in cases of incorrect responses.

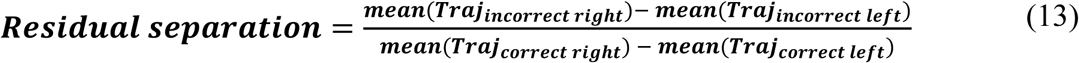

Mean (*Traj*) represents averaged trajectory at the time corresponding to each mode.

### Linear discriminant analysis

This procedure closely follows the previously described methodologies^51,52^. In investigating a neural network that processes stimulus-tuned input under noisy conditions, with the objective of accurately identifying the stimulus, we examined the characteristics of neural signals. Neural signals are assumed to be modeled as a combination of stimulus-driven signals and noise. The noise is characterized as temporally uncorrelated and independent of the stimulus. The analysis identifies a statistically optimal strategy for distinguishing between two different signals. This strategy involves performing a linear projection combined with temporal filtering of the input time series. By applying this approach, we derived the optimal projection weights and filtering mechanisms. Additionally, we calculated the signal-to-noise ratio (SNR) achieved through specific combinations of projection and filtering. The trials were categorized into four groups based on the mouse’s response to directional cues: (1) Trials where the mouse licked right in response to a right-directed cue. (2) Trials where the mouse licked left in response to a left-directed cue. (3) Trials where the mouse licked left in response to a right-directed cue. (4) Trials where the mouse licked right in response to a left-directed cue.

For the task of distinguishing between two types of signals (as an example, only one pair of response and cue listed above) *S*_*i*_, where *i* = 1 corresponds to a correct right decision and *i* = 2 to a correct left decision, we consider the dynamics of signals at each time step *t*. The assumption of input signal vector, ***u***(*s*_*i*_, *t*), follows a multivariate normal distribution, characterized by a stimulus-dependent mean and stimulus-independent covariances. The mean signal vector for the stimulation *S*_*i*_ is denoted as ***m***_*i*_ and total mean vector as ***m***. The within signal class covariance matrix is calculated as:

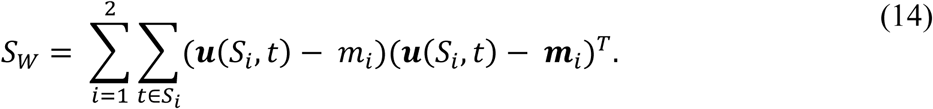

The covariance between signal class matrix is calculated as:

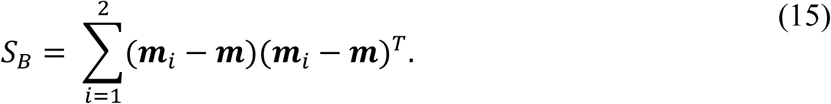

From these matrices, transformation matrix of LDA can be derived

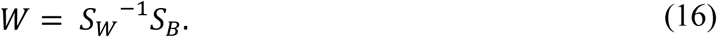

The eigen vector ***v***_*MAX*_ of maximal eigenvalue *λ*_*MAX*_ is selected as a maximum separation vector for discrimination of signal that we can project signal to calculate the SNR.

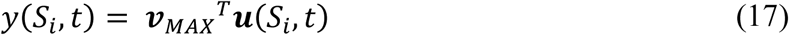

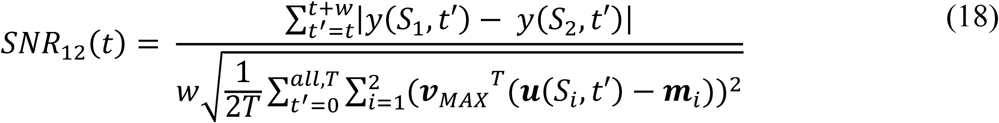

To standardize comparisons across different sessions and ensure a uniform basis for analysis, the SNR values observed were normalized against the maximum SNR value observed in the respective session. This step was critical for drawing direct comparisons across sessions, effectively illuminating the signal quality differences between them. The utilization of Matlab for all data analysis provided a robust framework for dissecting and understanding the dynamic processes underlying the behavioral responses.

### Declaration of generative AI and AI-assisted technologies in the writing process

During the preparation of this work, the authors used ChatGPT-OpenAI to correct grammatical errors. After using this tool, the authors reviewed and edited the content as needed and take full responsibility for the content of the publication.

## Acknowledgments

This work was supported by Korea Brain Research Institute Research Programs (grant 23-BR-01-01, 23-BR-03-01, 24-BR-03-08, and 23-BR-04-04 [JCR]) and National Research Foundation of Korea, under the Ministry of Science and Information and Communication Technology (grant NRF-2017M3A9G8084463 and NRF-2022R1A2C1004216 [JCR]).

## Author contributions

Conceptualization-J.C.R, J.H.C, S.W.B; Data curation-S.W.B, J.H.C, M.S.Y; Formal analysis-J.H.C, S.W.B, M.S.Y; Funding acquisition-J.C.R; Investigation-J.H.C; Methodology-J.C.R, J.H.C, J.H.P, S.W.B; Project administration-J.C.R; Resource-J.C.R; Supervision-J.C.R; Validation-J.C.R, L.I.S, C.H.K, J.W.C; Visualization-S.W.B, M.S.Y; Writing-J.C.R, S.W.B; Writing-review & editing-J.C.R, S.W.B;

## Supplementary information

**Supplementary Figure 1. Time-correlation of activity dynamics along the coding direction by the PPC neurons in the STM task.**

**A.** Time-wise Pearson’s correlation of activity vectors for direction-selective posterior parietal cortex (PPC) neurons during the short-term memory (STM) task. Direction Selectivity was determined using a modified Generalized Linear Model (GLM) as described in the Methods section. Neural activity was binned over 0.5-s intervals across 11 sessions from 6 mice (total neurons: n=2,437).

**B.** Time-correlation analysis for non-direction-selective neurons (neurons with *p* > 0.01): Neural activity was similarly binned over 0.5-s intervals across the same 11 sessions from 6 mice (total neurons: n=2,319).

**Supplementary Figure 2. Logic diagram of directional and non-directional phase-selective neurons identified by the modified General Linear Model (GLM).**

**A.** Fraction of sample-phase specific neurons showing significant response selectivity to left or right visual stimuli. The overlap between the circles represents the neurons that responded to both directions.

**B.** Same as A but for delay-phase specific neurons.

**C.** Same as A but for reward-phase specific neurons.

**Supplementary Figure 3. Reliability index of directional and non-directional neurons.**

**A.** Reliability index (RI) histogram for direction-selective neurons in correct trials (blue) and shuffled trials (gray). PD, preferred direction; OD, opposite direction.

**B.** RI histogram for non-directional neurons in correct trials (orange) and shuffled trials (gray). L, left direction; R, right direction.

**C.** Cumulative distribution function (CDF) of RI for directional single-phase selective neurons (blue) and non-directional single-phase selective neurons (orange). The Kolmogorov-Smirnov (KS) test revealed a significant difference between the two distributions (D = 0.3791, *p* = 5.5476e-71).

**D.** RI histogram of directional, sample-phase neurons in correct trials (blue) and shuffled trials (gray).

**E.** RI histogram for non-directional, sample-phase selective neurons in correct trials (orange) and shuffled trials (gray).

**F.** CDF of RI for directionally selective (blue) and non-directionally selective (orange) neurons during the sample phase. The K-S test revealed a significant difference between the distributions (D = 0.3689, *p* = 5.0852e-29).

**G.** Same as panel D, but for directional, delay-phase selective neurons.

**H.** Same as panel E, but for non-directional, delay-phase selective neurons.

**I.** Same as panel F, but for delay-phase selective neurons. The K-S test revealed a significant difference between the distributions (D = 0.3783, *p* = 4.1962e-13).

**Supplementary Figure 4. Phase selectivity of GLM-defined phase-selective and non-selective neurons.**

**A.** Phase selectivity index (PSI) histogram for sample-phase selective neurons (black) and non-phase selective neurons (gray) in correct trials. A more positive PSI indicates a stronger response specificity to the corresponding phase. S, sample phase; D, delay phase; R, reward phase.

**B.** PSI histogram for delay-phase selective neurons (blue) and non-phase selective neurons (gray) in correct trials.

**C.** PSI histogram for reward-phase selective neurons (red) and non-phase selective neurons (gray) in correct trials.

**Supplementary Figure 5. Anatomical distances and functional similarities.**

**A.** The relationship between within-group and across-group distances for phase-selective neurons.

**B.** The within-group and across-group distances for directional and non-directional neurons.

**Supplementary Figure 6. Reliability index of direction- and phase-selective neurons in error trials.**

**A.** Color-coded responses of non-directional single-phase selective neurons in error trials. Trial-averaged ΔF/F was normalized to the maximum ΔF/F in correct trials and aligned based on the time-to-peak of each neuron.

**B.** Preference index (PI) histogram of GLM-defined non-directional single-phase selective neurons in correct (orange), error (red open), and shuffled (gray) trials.

**C.** PI histogram of non-directional sample-phase selective neurons in correct (black), error (red open), and shuffled (gray) trials.

**D.** PI histogram of non-directional delay-phase selective neurons in correct (blue), error (red open), and shuffled (gray) trials.

**E.** Histograms of reliability indices (RI) of non-directional single-phase selective neurons in correct (orange), error (red open), and shuffled (gray) trials.

**F.** RI histograms of non-directional sample-phase selective neurons in correct (black), error (red open), and shuffled (gray) trials.

**G.** RI histograms of non-directional delay-phase selective neurons in correct (blue), error (red open), and shuffled (gray) trials.

**H.** Relationship between the RI difference in error trials and the initial RI in correct trials. Δ (Reliability index) is defined as the difference in RI in error trials (RI_incorrect_ – RI_correct_).

**I.** Same scatter plot as in H but for non-directional sample-phase selective neurons.

**J.** Same scatter plot as in H but for non-directional delay-phase selective neurons within the posterior parietal cortex (PPC).

**Supplementary Figure 7. Variance explained by the number of principal components.**

**A.** The fraction of variance explained (bar) and the accumulated variance explained (line) as principal components of right-directional phase-specific neurons are added.

**B.** Same as in A, but for left-directional phase-specific neurons.

**C.** Same as in A, but for combined directional-selective neurons

**Supplementary Figure 8. Discriminability between the trajectory distances and schematic of the trajectory similarity index (TSI).**

**A.** Area under the receiver operating characteristics curve (auROC) between d1 and d2 (blue) and between d1 and d5 (red) in Fig. 5 over time. ROC analysis was conducted for the activity distances over time. A greater absolute value indicates better separation. For calculating trajectory distances (d1-d5), 50% of the neurons were used to create a vector, and the remaining 50% were projected onto it. This process was repeated over 100 random trials.

**B.** auROC between d0 and d3 (blue) and between d0 and d4. Note that values closer to -1 indicate better separation.

**C.** At each time point, the trajectory distance from the trial-averaged values to the mean correct preferred signal trial (PP) and the distance to the correct opposite signal trial (OO) were measured to calculate the trajectory selectivity index (TSI), as shown in the formula. If the measured population activity at a given time is similar to the PP response, the TSI will be positive. Conversely, activities similar to the OO response will result in a negative TSI value.

**Supplementary Figure 9. Framework of the proposed method for automatic event-related region of interest (ROI) detection.**

**A.** The proposed method detects an event-related neuron pixelwise.

**B.** Pixelwise error optimization with amplitude of calcium signal β.

**C.** The 1-sampled *t*-test is applied to the β distribution of each pixel.

**D.** Selected ROI pixels are clustered automatically.

