## Supplementary material for "Short-term memory errors are strongly associated with a drift in neural activity in the posterior parietal cortex": Supplemetary figure

### Supplementary Figure

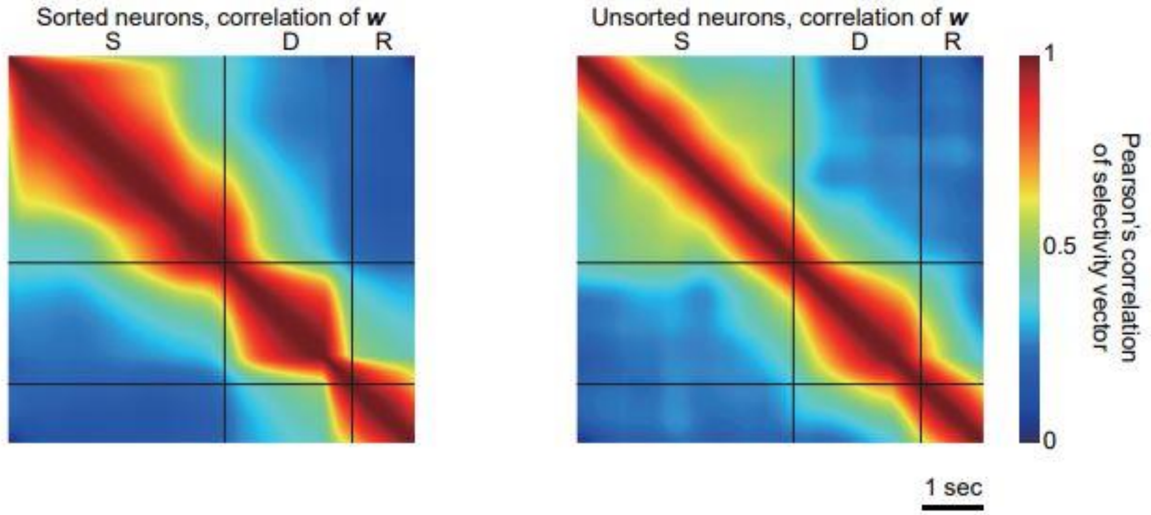


**Supplementary Figure 1. Time-correlation of activity dynamics along the coding direction by the PPC neurons in the STM task. A.** Time-wise Pearson's correlation of the activity vector by the direction-selective PPC neurons. Direction selectivity was determined by the modified Generalized Linear Model (GLM) described in the Methods section. Activity was binned over 0.5 seconds across 11 sessions from 6 mice (n = 2,437). **B.** Time-correlation of neurons that did not show significant directional selectivity ($p > 0.01$). Activity was binned over 0.5 seconds across 11 sessions from 6 mice (n = 2,319).

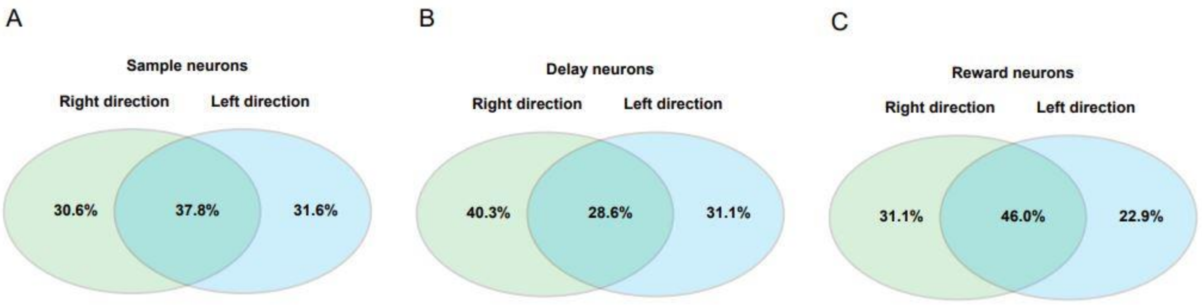


**Supplementary Figure 2. Logic diagram of directional and non-directional phase-selective neurons identified by the modified General Linear Model (GLM). A.** Fraction of sample-phase specific neurons showing significant response selectivity to left or right visual stimuli. The overlap between the circles represents the neurons that responded to both directions. **B.** Same as A but for delay-phase specific neurons. **C.** Same as A but for reward-phase specific neurons.

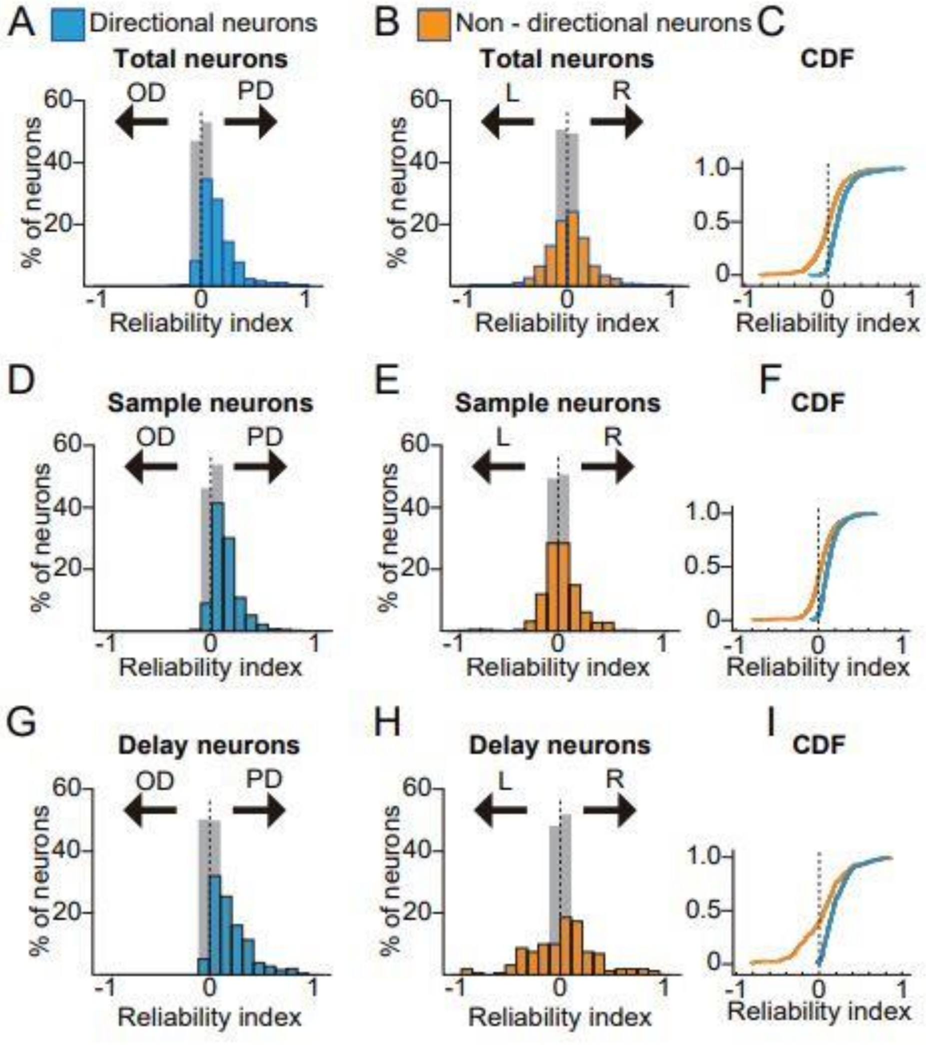


**Supplementary Figure 3. Reliability index of directional and non-directional neurons.**

**A.** Reliability index (RI) histogram of direction-selective neurons in correct trials (blue) and shuffled trials (gray). PD, preferred direction; OD, opposite to preferred direction. **B.** RI histogram of non-directional neurons in correct (orange) and shuffled (gray) trials. L, left direction; R, right direction. **C.** Cumulative distribution function (CDF) of RI for directional single-phase specific neurons (blue) and non-directional single-phase specific neurons (orange). The Kolmogorov-Smirnov (KS) test revealed a significant difference between the two distributions (D = 0.3791, p = 5.5476e-71). **D.** RI histogram of directional, sample-phase neurons in correct trials (blue) and shuffled trials (gray). **E.** RI histogram of non-directional selective, sample-phase neurons in correct (orange) and shuffled (gray) trials. **F.** CDF of RI for directionally selective (blue) and non-selective (orange) neurons. The K-S test revealed D = 0.3689, p = 5.0852e-29. **G.** Same as D but for delay-phase selective, directional neurons. **H.** Same as E but for delay-phase selective, directional neurons. **I.** Same as F but for delay-phase selective, non-directional neurons. The K-S test revealed D = 0.3783, p = 4.1962e-13.

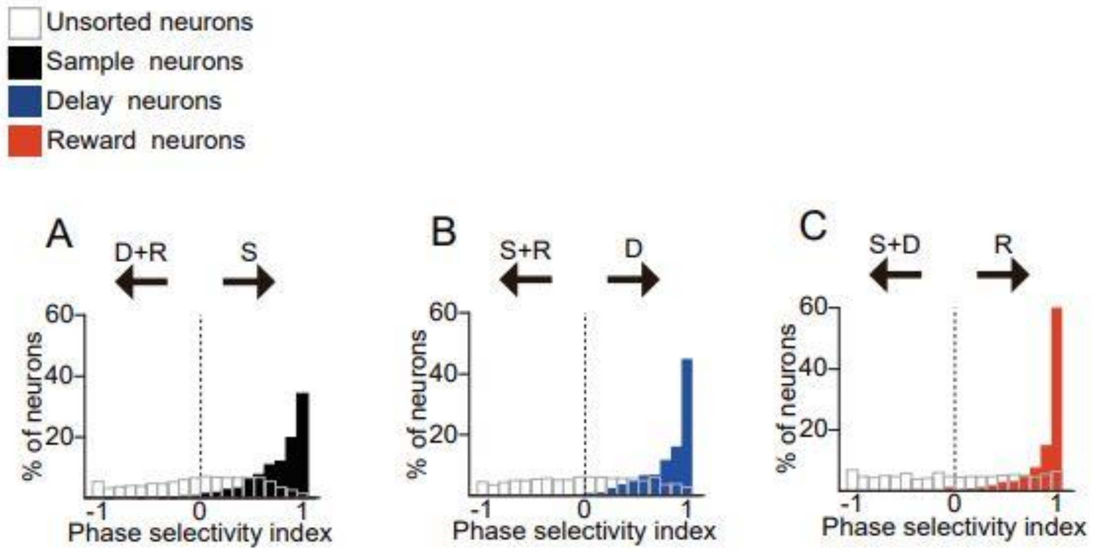


**Supplementary Figure 4. Phase selectivity of GLM-defined phase-selective and non-selective neurons. A.** Phase selectivity index (PSI) histogram of sample-phase selective neurons in correct trials (black) and non-phase selective neurons in correct trials (gray). A more positive PSI indicates a more specific response to the corresponding phase. S, sample phase; D, delay phase; R, reward phase. **B.** PSI histogram of delay-phase neurons (blue) and non-phase selective neurons (gray) in correct trials. **C.** PSI histogram of reward-phase neurons (red) and non-phase selective neurons (gray) in correct trials.

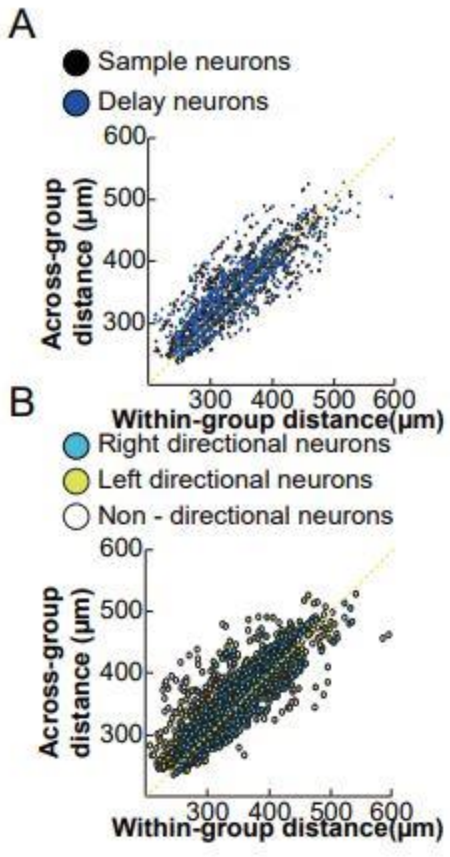


**Supplementary Figure 5**. **Anatomical distances and functional similarities. A.** The relationship between within-group and across-group distances for phase-selective neurons. **B.** The within-group and across-group distances for directional and non-directional neurons.

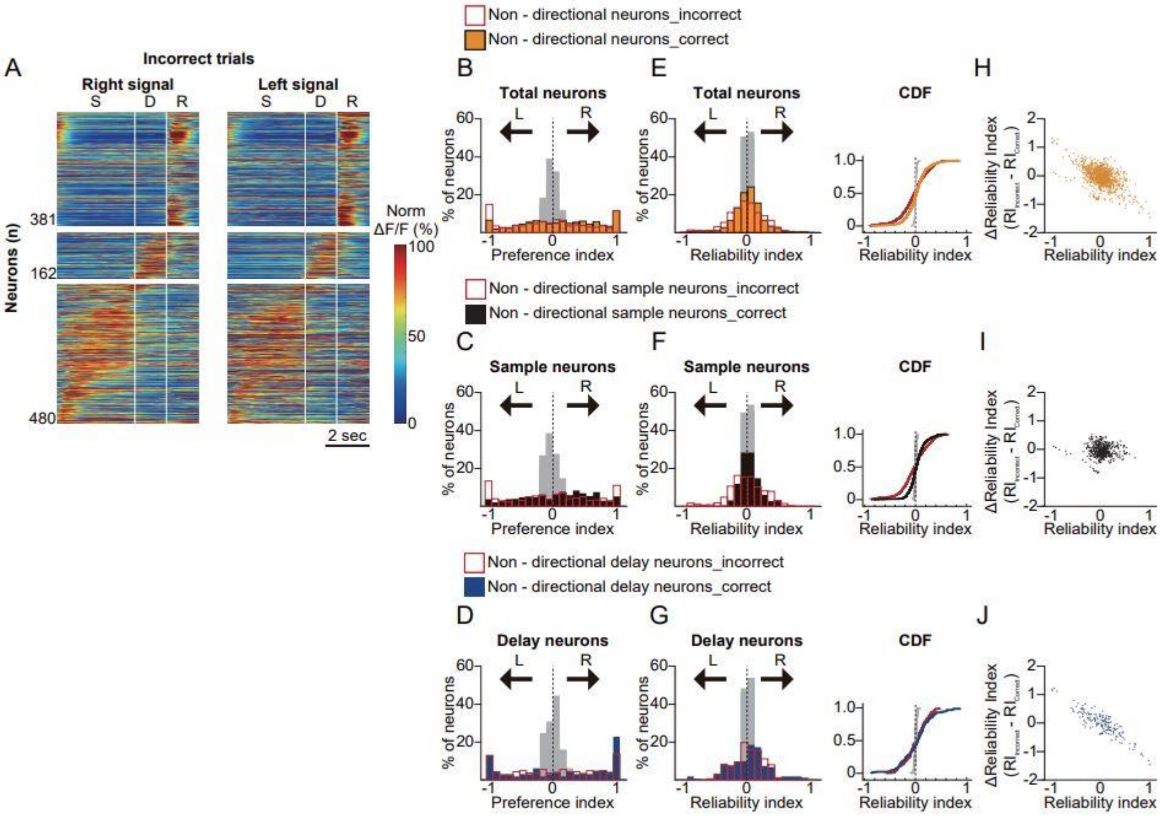


**Supplementary Figure 6. Reliability index of direction- and phase-selective neurons in error trials. A.** Color-coded responses of non-directional single-phase selective neurons in error trials. Trial-averaged ΔF/F was normalized to the maximum ΔF/F in correct trials and aligned based on the time-to-peak of each neuron. **B.** Preference index (PI) histogram of GLM-defined non-directional single-phase selective neurons in correct (orange), error (red open), and shuffled (gray) trials. **C.** PI histogram of non-directional sample-phase selective neurons in correct (black), error (red open), and shuffled (gray) trials. **D.** PI histogram of non-directional delay-phase selective neurons in correct (blue), error (red open), and shuffled (gray) trials. **E.** Histograms of reliability indices (RI) of non-directional single-phase selective neurons in correct (orange), error (red open), and shuffled (gray) trials. **F.** RI histograms of non-directional sample-phase selective neurons in correct (black), error (red open), and shuffled (gray) trials. **G**. RI histograms of non-directional delay-phase selective neurons in correct (blue), error (red open), and shuffled (gray) trials. **H.** Relationship between the RI difference in error trials and the initial RI in correct trials. Δ (Reliability index) was defined as RI changes in error trials ($RI_{incorrect}$ - $RI_{correct}$). **I.** Same scatter plot as in H but for non-directional sample-phase selective neurons. **J.** Same scatter plot as in H but for non-directional delay-phase selective neurons within the PPC.

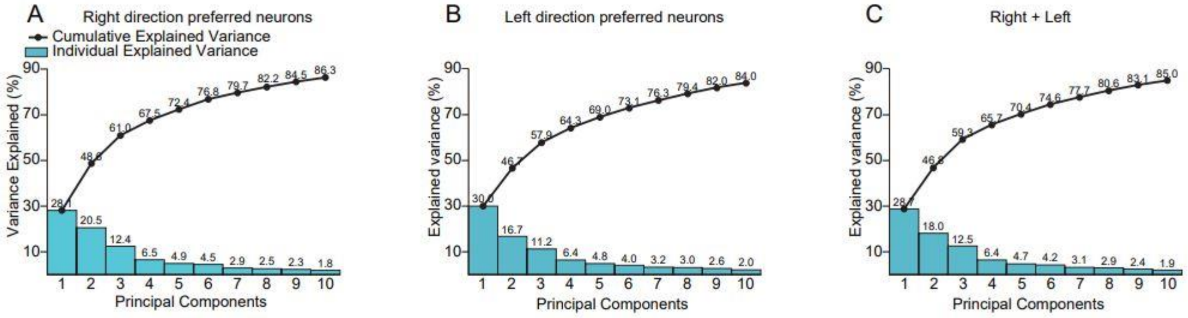


**Supplementary Figure 7. Variance explained by the number of principal components. A.** The fraction of variance explained (bar) and the accumulated variance explained (line) as the principal components of right-directional phase-specific neurons are added. **B.** Same as in A, but for left-directional phase-specific neurons. **C.** Same as in A, but for combined directional-selective neurons

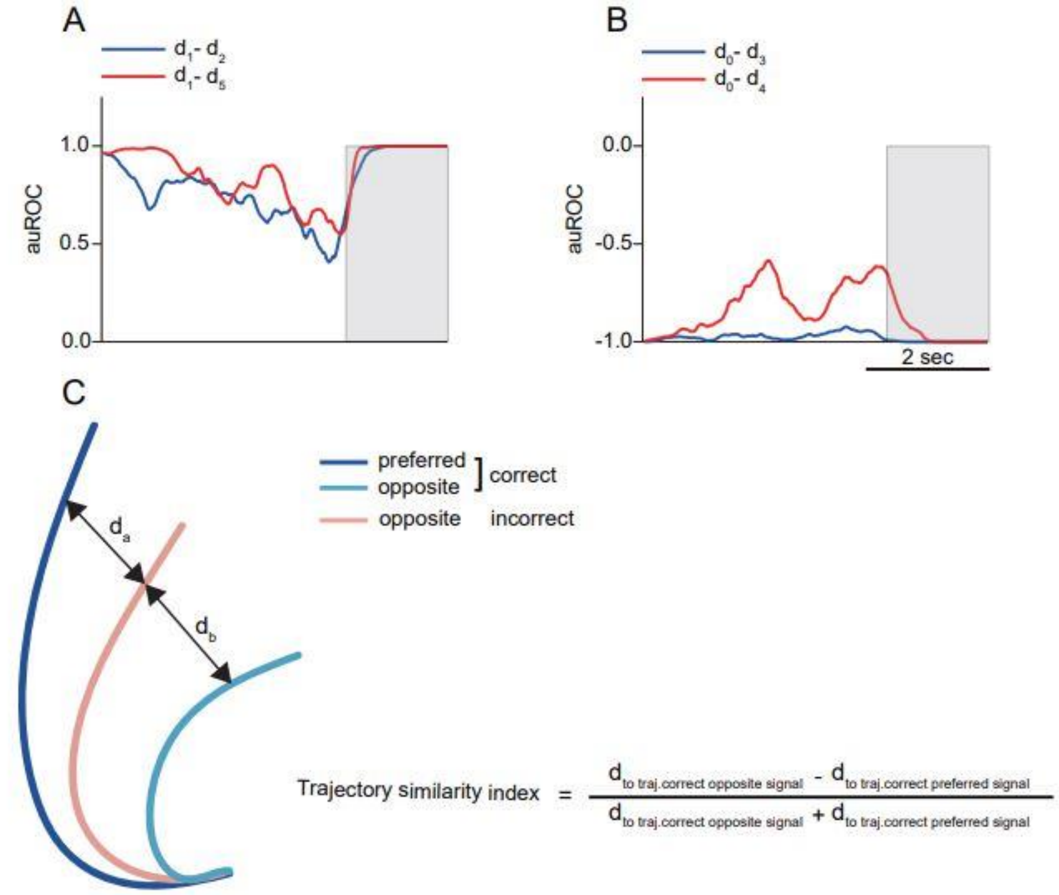


**Supplementary Figure 8. Discriminability between the trajectory distances and schematic of the trajectory similarity index (TSI). A.** Area under the receiver operating characteristics curve (auROC) between d1 and d2 (blue) and between d1 and d5 (red) in Fig. 5 over time. ROC analysis was conducted for the activity distances over time. A greater absolute value indicates better separation. When calculating trajectory distances (d1-d5), 50% of the neurons were used to create a vector, and the remaining 50% were projected onto it. This process was repeated over 100 random trials. **B.** auROC between d0 and d3 (blue) and between d0 and d4. Note that values closer to -1 indicate better separation. **C.** At each time point, the trajectory distance from the trial-averaged values to the mean correct preferred signal trial (PP) and the distance to the correct opposite signal trial (OO) were measured to calculate the trajectory selectivity index (TSI), as shown in the formula. If the measured population activity at a given time is similar to the PP response, the TSI will be positive. Conversely, activities similar to the OO response will result in a negative TSI value.

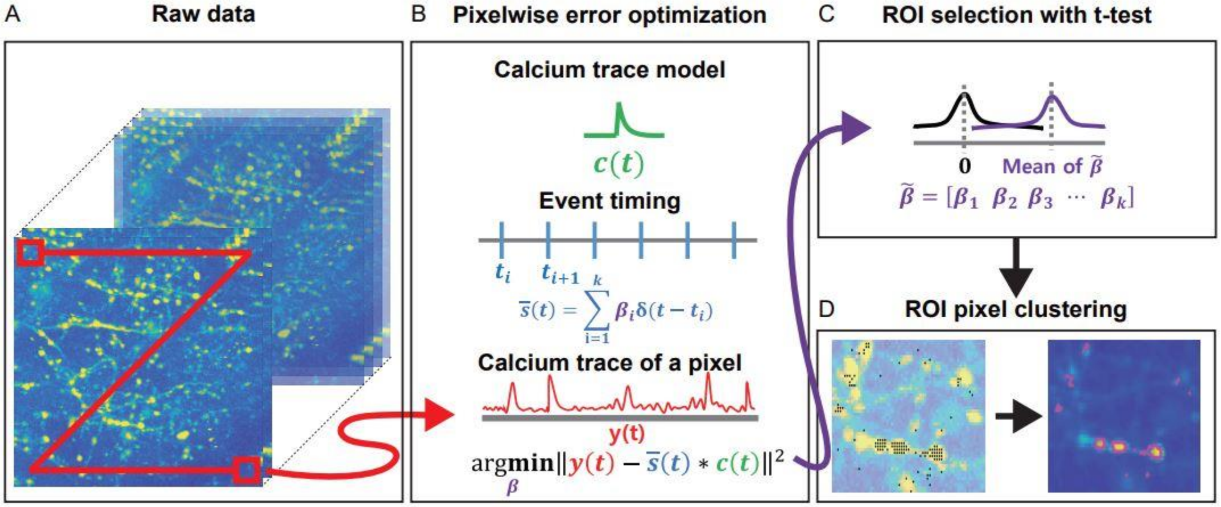


**Supplementary Figure 9. Framework of the proposed method for automatic event-related ROI detection. (A)** The proposed method detects an event-related neuron pixelwise. **(B)** Pixelwise error optimization with amplitude of calcium signal β. **(C)** The 1-sampled t-test is applied to the β distribution of each pixel. **(D)** Selected ROI pixels are clustered automatically.
